# An Efficient CRISPR protocol for generating Conditional and Knock-in mice using long single-stranded DNA donors

**DOI:** 10.1101/141424

**Authors:** Hiromi Miura, Rolen M. Quadros, Channabasavaiah B. Gurumurthy, Masato Ohtsuka

## Abstract

The CRISPR/Cas9 tool can easily generate knockout mouse models by disrupting the gene sequence, but its efficiency for creating models that require either insertion of exogenous DNA (knock-in) or replacement of genomic segments is very poor. The majority of mouse models used in research are knock-in (reporters or recombinases) or gene-replacement (for example, conditional knockout alleles containing *LoxP* sites flanked exons). A few methods for creating such models are reported using double-stranded DNA as donors, but their efficiency is typically 1–10% and therefore not suitable for routine use. We recently demonstrated that long single-stranded DNAs serve as very efficient donors, both for insertion and for gene replacement. We call this method *Easi*-CRISPR (efficient additions with ssDNA inserts-CRISPR), a highly efficient technology (typically 25%-50%, and up to 100% in some cases), one that has worked at over a dozen loci thus far. Here, we provide detailed protocols for *Easi*-CRISPR.

## INTRODUCTION

The latest advances in genome editing technologies, specifically those using the CRISPR/Cas9 system, have helped simplify the process of generating genetically engineered animal models. Genome editing is performed, in cells or in zygotes, through two molecular events. First, the Cas9 nuclease is taken to the target site through a guide RNA and the nuclease causes a double-stranded DNA break (DSB) at the target site. In the second step, the DSB is joined back by one of the two major DSB repair events, either through: (a) non-homologous end joining (NHEJ), which usually leads to a change in the nucleotide sequence; or (b) homology-directed repair (HDR), if an exogenous repair template is supplied that contains homology arms.

During the past 3–4 years, CRISPR/Cas9 technology has significantly impacted how mouse genome engineering is performed.^1,2^ It is now used routinely to generate gene disruptions through short insertions or deletions (*indels*) via NHEJ, and also to insert short exogenous sequences provided as single-stranded oligodeoxynucleotides (ssODNs) via HDR. The ssODN repair templates used are typically 100–200 bases long, consisting of a few bases of altered sequence flanked by homology arms of 40–80 bases.^3,4^ In contrast, double-stranded DNAs (dsDNAs) are used as repair templates for projects requiring insertion of longer sequences (such as reporter/recombinase knock-ins). Compared with ssODN donors, however, the insertion efficiency of dsDNA donors is poor,^5^ often requiring homology arms of at least 0.5–1 kb or longer.^6,7^ Creating conditional knockout models requires even higher technical perfection because these involve replacing a gene fragment (target exon/s) with a floxed (*LoxP* flanked exon) cassette gene replacement.

### Development of Easi-CRISPR

We hypothesized that if longer ssDNAs can be used as donors, they, too, would be inserted at a similar efficiency as ssODNs, typically higher than that of the dsDNA donors. When we began to test our hypothesis, single-stranded DNAs (ssDNAs) of longer than 200 bases were not commercially synthesizable. To generate longer ssDNAs, we used classical molecular biology steps such as converting dsDNA to an RNA (using an *in vitro* transcription step) and reverting the RNA back to DNA (using a reverse transcriptase step) to obtain ssDNAs. We named this strategy (synthesizing ssDNAs from dsDNA templates) *“in vitro* transcription and reverse transcription,” or *iv*TRT. Using long ssDNAs as donor templates, we developed highly efficient CRISPR-HDR protocols in three stages, as described below.

In the *first stage*, we demonstrated that longer sequences of ~400 bases of new sequences with ~55 bases homology arms on each side could be inserted at Cas9 cleavage sites.^8^ These experiments demonstrated that insertion of artificial microRNA sequences into an intronic site of *eEF2* locus resulted in knock-down of target genes (eGFP and *Otx2*) as analyzed at the embryonic stages. Insertion efficiencies were up to 83% of offspring.

Extending this work further, in the *second stage*, we demonstrated that sequences up to ~1.5 kb can be inserted to express various types of proteins^9^. Our aim in those experiments was to further develop the method to achieve generation of germ-line, transmittable knock-in models that can express protein coding sequences such as recombinases, reporters, or transcriptional inducers. Indeed, the method worked at very high efficiency for precisely fusing expression cassettes to specific codons of genes. The method is shown to be robust (consistent performance at over six loci) and is highly efficient (25% to 67%).

As the *third stage* of developing high efficiency HDR protocols, we tested whether a gene segment can be excised out by making two cleavages in the genome, and replacing the segment with a modified piece (such as a floxed-exon cassette). The method worked very efficiently at many loci (8.5–100%).

The sgRNA and Cas9 mRNA were used as CRISPR reagents composition in the first stage of experiments, whereas separated crRNA + tracrRNA and Cas9 Protein (ctRNP) were used in the second and third stage experiments. ^9,10^

Systematic testing of the above three ssDNA insertion approaches led to our establishing streamlined protocols for generating many types of commonly used mouse models, including knock-down, knock-in, and conditional knockout mouse models. These protocols have worked at high efficiency for over a dozen loci, and experiments have been performed in at least three different laboratories^9^. We named the method *Easi*-CRISPR (<u>*E*</u>fficient <u>a</u>dditions with <u>s</u>sDNA inserts), and here we provide detailed protocols for Easi-CRISPR.

### Applications of the <u>Easi</u>-CRISPR method

Many types of genetically engineered mouse (GEM) models are used in biomedical research, and most of these rely on precise insertion of donor cassettes. Examples of a few commonly used models are: (i) floxed models that contain two *LoxP* sites flanking a gene segment to allow for conditional deletion of the gene; (ii) Cre or CreER^T2^ driver lines that are used to delete a floxed gene in a given tissue and/or at a given time; (iii) reporter strains used to monitor Cre specificity and sensitivity and/or to monitor gene expression; and (iv) inducible transcriptional regulator strains (rtTA/tTA) that permit doxycycline-regulated expression of tetO-containing promoters. The *Easi*-CRISPR approach can be used for generating all of the above types of animal models. In addition, because the *Easi*-CRISPR strategy works very efficiently for gene replacements (demonstrated up to about 1 kb ssDNAs thus far), the method can also be used for developing multiple point mutation knock-in animal models, and for swapping gene segments from other species (e.g., humanized animal models) in this size range. The method is also applicable to generate knock-down mice by introducing an artificial microRNA sequence at the intronic region of an endogenous gene, as described in Miura et al. (2015).^8^

This long ssDNA-based knock-in method was developed using micro-injection delivery in mice. The approach can also be adapted for use with electroporation or hydrodynamics gene delivery.^11,12^ *Easi*-CRISPR can also be used to generate genome-edited animals in other species wherever zygote/embryo delivery of CRISPR components (microinjection- or electroporation-based) is possible (other model organisms such as fly, zebrafish, rat, guinea pig, and rabbit, as well as livestock species such as cattle, sheep, goats and pigs).

### Comparison of other methods of knock-in and conditional knockout model generation

Mouse genome engineering has been broadly performed over three decades for two purposes: 1) direct injection of DNA to create transgenic models where DNA is integrated at random genomic locations; and 2) to create knock-in and (conditional) knockout models using classical homologous recombination-mediated gene targeting in embryonic stem (ES) cells. The ES cell-based methods were the only choices before programmable nucleases were demonstrated for gene targeting. However, those methods are laborious, time-consuming, expensive, and, more importantly, may not lead to generation of a germ line-established mutant line. Soon after the CRISPR/Cas system was developed for genome editing in mammalian cells,^13,14^ it was demonstrated that mouse knock-in and conditional knockout alleles could be generated rapidly.^6^ The research community anticipated that this technology would soon replace ES cell-based gene targeting,^15^ because the required components can be introduced directly into mouse embryos using the same microinjection techniques commonly used for random transgenesis. However, many laboratories have been unsuccessful in employing CRISPR strategies for generating knock-in and conditional knockout alleles.^16^

One of the main reasons for this lack of success is that HDR efficiency using dsDNA donors is generally very poor. However, a few strategies for increasing the targeting efficiency of dsDNA donors have been reported. These include: (1) inhibition of NHEJ or enhancement of HDR through chemical treatments.^17,18^ Such methods have not become popular because they only provide a marginal gain in their efficiencies, and these strategies have been shown to be toxic to cells, disrupting fundamental DNA repair processes.^19^ (2) Certain strategies use circular donors with built-in synthetic guide sequences. The donors are linearized inside the cell/embryo through Cas9 cleaving at the synthetic guide recognition sites, to enhance their insertion at the target site. Although these latter strategies offer better alternatives to those that perturb DNA repair, and have efficiencies reaching up to 37.5%,^20^ they too have limitations, because custom donor plasmids must be constructed for each target site.

In direct comparison to circular dsDNA donor-based strategies, *Easi*-CRISPR offers several advantages: (1) donor DNAs are simple to design and construct with no need for long homology arms (~50–100 base long arms are enough); (2) efficiency is very high (typically 25–75%, reaches up to 100% for some loci), and (3) the method works for many loci. The linear dsDNAs are the standard forms of DNA cassettes used in generating transgenic animals (where the DNA is inserted at random locations). Although injection of dsDNAs would lead to many events of random transgene insertions, in our first-stage experiments developing *Easi*-CRISPR, we compared targeted insertion efficiency by dsDNA (PCR product) vs ssDNA (synthesized using ivTRT) with the same sequence (and the same lengths of homology arms). We found that ssDNAs had more than two times higher efficiency than dsDNAs.^8^ We also found that the viability of injected embryos was much higher when ssDNA was used as a repair template compared to dsDNA, and that this was probably due to dsDNA toxicity.

Conditional knock-out models are the most highly produced type of genetically engineered models. The genetic manipulation required to create such alleles using CRISPR/cas strategies involves a much higher level of technical perfection than for knock-in designs because it requires two cleavages in the genome and precise replacement of the gene segment with a floxed DNA cassette between the cleavage sites. An approach where two ssODN donors could be used to insert *LoxP* sites, involving two cleavages in the genome, was initially feasible,^6^ but this approach has been considered highly challenging due to many undesired outcomes. These include insertion of only one *LoxP*, deletion of the target exon between the genomic cleavages, and insertion of *LoxP* sites *in trans* ^21^. The method has also failed otherwise for many loci.^16^ Easi-CRISPR is less likely to result in these unwanted outcomes, especially compared to the two-ssODN oligo-based method. Additionally, using circular dsDNAs (as standard knock-in donors) and inserting them through nicking (using Cas9 nickase) has been reported for generating conditional alleles,^22^ but this approach is also not used routinely due to low efficiency.

## Experimental Design

The *Easi*-CRISPR method involves four major steps: (1) designing of an *Easi*-CRISPR strategy; (2) synthesis and purification of ssDNA and other CRISPR components for microinjection; (3) preparation of *Easi*-CRISPR components and their microinjection into mouse zygotes; and (4) genotyping of offspring. A diagram of the workflow is presented in **Figure 1**.

**Figure 1:**
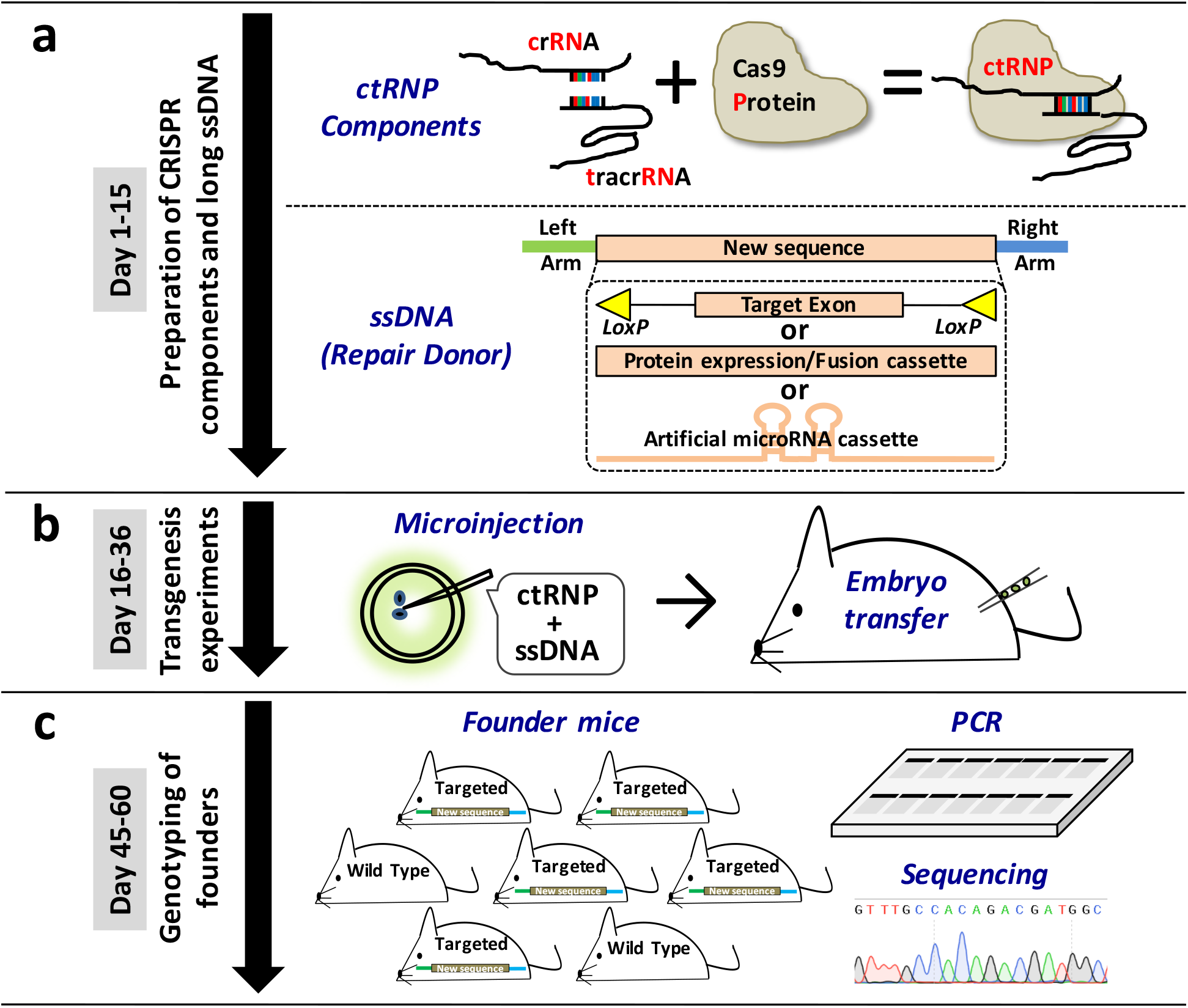
Schematic of *Easi*-CRISPR. The procedure involves three broad steps: **(a)** Assembling of CRISPR Ribonucleoprotein components (crRNA + tracrRNA + Cas9 Protein: ctRNP) and generating a long ssDNA donor. **(b)** Preparation of *Easi*-CRISPR components, their microinjection into mouse zygotes and generation of founder offspring, **(c)** Genotyping of offspring.

### Designing of an *Easi*-CRISPR strategy

The strategy involves two broad steps: searching for CRISPR targets, and designing ssDNA repair cassettes

### Searching for CRISPR targets

Guide search is a very common step in CRISPR genome engineering designs, and has been described in numerous protocols including mouse genome engineering^23,24^. Genomic sequence around the region of interest is retrieved from a genome browser such as http://asia.ensembl.org/index.html and the CRISPR target sites using CHOPCHOP (http://chopchop.cbu.uib.no/index.php) and/or CRISPR Design (http://crispr.mit.edu/) sites. Select the site as close as possible to the insertion site.

The guide selection principles relevant to different applications of *Easi*-CRISPR are described in **Box 1**

#### Box 1: Principles of guide selection for *Easi*-CRISPR applications.

The *Easi*-CRISPR method was developed to be suitable for most commonly used mouse model designs, including: (a) insertion of protein coding sequences to fuse with genes (also known as knock-in models); (b) conditional knockout mouse models (also known as floxed models); and (c) knocking-down genes through insertion of artificial microRNA sequences into intronic sites. In all three designs, long ssDNAs are used as donors, in contrast to the differing location of guides on genes and the types of insertion cassettes among designs. The principles of guide selection for these designs are described here:

##### Floxing

Determine the target exon, the deletion of which will lead to predicted to loss of protein expression upon Cre-mediated recombination. *LoxP* sites are inserted at the flanking introns to the target exon, and then chosen within the flanking introns to the target exon. Because it is difficult to accurately predict the regulatory sequences surrounding the exons necessary for splicing (e.g. splicing donor/acceptor sites, branch site, and polypyrimidine tract), we suggest placing *LoxP* sites at least 100 bases away from the intron/exon boundaries. Care should be taken not to insert *LoxPs* at the evolutionary conserved regions to avoid disruption of functional sequences such as enhancers (see below).

##### Knocking in (or fusing) coding sequences with genes

To determine the targeted site, using either 5’ end or 3’ end of the gene (start or stop codons respectively). Identify guides that recognize sequences very close to the start or stop codons for precise fusion.

##### Inserting artificial microRNA cassettes for knocking down genes

Our first proof of principle work (that long ssDNAs serve as efficient donors) demonstrated that artificial microRNA sequences can be inserted at introns of genes (host gene). Inserted cassettes are transcribed as part of the host gene transcription and eventually processed into mature microRNAs to knockdown the protein expression of their target genes. We tested introns of the *eEF2* gene, which enabled ubiquitous expression of microRNA sequences. More such sites in other ubiquitously expressing genes and/or introns of genes that will have desired tissue-specific expression patterns can be explored for inserting artificial microRNAs. A few designing principles for this application: 1) Determine the targeted intron to be inserted. 2) Avoid choosing evolutionary conserved regions that potentially contain sequences of some important (albeit unknown) function; choosing such regions may inadvertently cause disruption of the evolutionarily functional sequences. 3) Similar to floxing design, choose regions sufficiently far away from exon boundaries to avoid disrupting splicing signals. Potential splicing events in a given sequence can be predicted using the GENESCAN tool (exon prediction algorithm; http://genes.mit.edu/GENSCAN.html). Both wild type (untargeted) and theoretical sequence of the modified locus (after cassette insertion) should be analyzed using this tool.

### Designing ssDNA repair cassettes

The typical architecture of cassettes constitutes two homology arms (left and right) and a middle region (new sequence to be inserted). Homology arms are about 55 to 100 bases long. A T7 RNA polymerase promoter site is included, just upstream of one of the homology arms (referred as proximal arm), and is needed for *in vitro* transcription. A unique restriction site is included after the distal arm (one that does not need a T7 promoter), to use it for linearizing the plasmid. The central region can be up to about 2 kb or more. The design principles and architecture of ssDNA donors and dsDNA templates for knock-in, floxing, and knock-down are shown in **Figures 2 to 4**. Note that the design principles for inserting artificial microRNA cassettes share some features between floxing and knock-in designs, involving one cleavage insertion (as in the case of knock-in), and insertion of cassettes at an intronic region (as in the case of floxing). The repair cassette should be carefully designed to not include the guide recognition sequence, and thus avoid re-cleaving by the Cas9/gRNA after the correct insertion.

**Figure 2:**
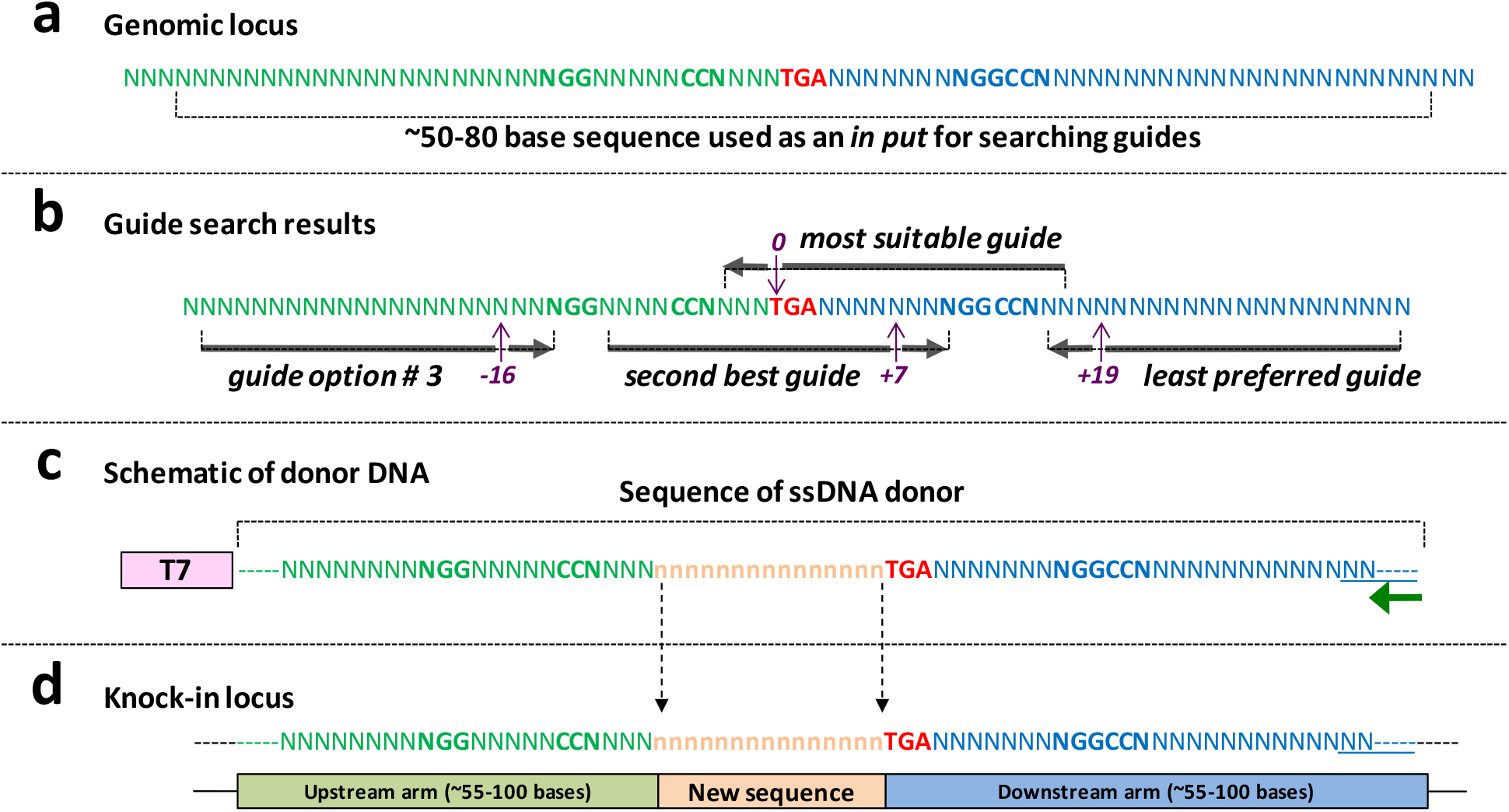
Design principles of knocking-in using *Easi*-CRISPR and the architecture of ssDNA donor. **(a)** Genomic locus of a hypothetical gene’s last exon showing the stop codon in red (TGA). The green sequence upstream of- and the blue sequence downstream of-stop codon will be parts of upstream and downstream arms of the ssDNA donor respectively. **(b)** Hypothetical guide search results showing four guide options with protospacer adjacent motif (PAM) sequences (5′-NGG-3′). The guide that cleaves immediately upstream of the stop codon will be the most suitable guide for *Easi*-CRISPR. If such a guide is not available, a next guide immediately close to the target site should be chosen. In the example shown here, a guide that cuts 7 bases downstream should be the second option. The other two guides (options 3 and the least preferred guide) cleave at −16 bases and +19 bases from the target site. Farther the guide form the target site, poorer will be the correct insertion frequency because imprecise insertion rates become higher. If either of the last two guides are chosen, the donor cassette should contain mutation/s in the guide recognition sites (or PAM) to prevent Cas9 re-cleaving after the cassette is inserted. **(c)** Schematic of donor DNA showing T7 promoter (as part of dsDNA template) and the actual part of ssDNA donor. T7 promoter sequence should be included in the dsDNA template (used for *iv*TRT). Green arrow shows the primer for reverse transcription. **(d)** Knock-in locus showing correct fusion of new sequence.

**Figure 3:**
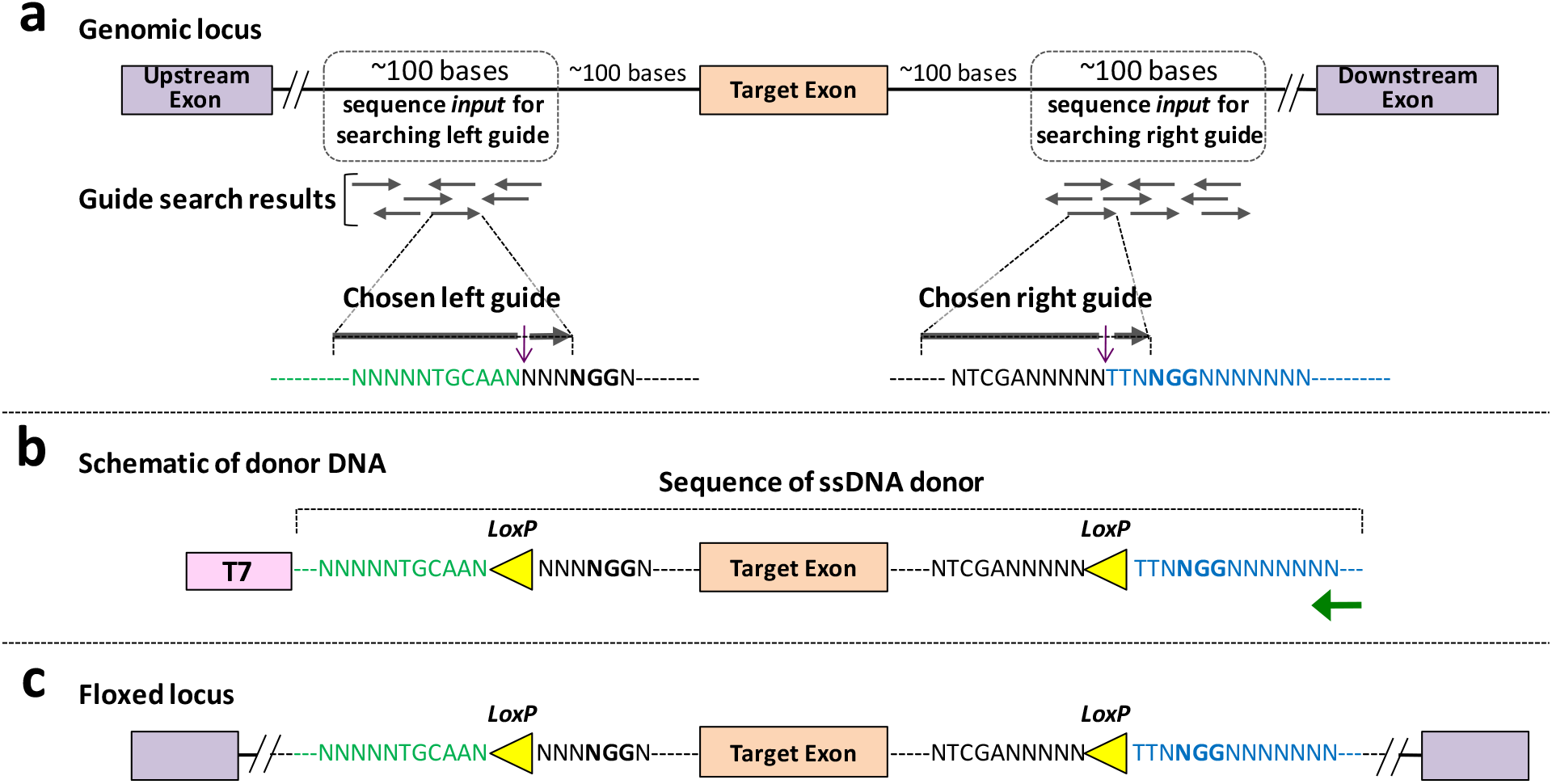
Design principles of floxing using *Easi*-CRISPR and the architecture of ssDNA donor. **(a)** Genomic locus of a hypothetical gene’s target exon and its surrounding regions chosen for guide search. Hypothetical guide search results showing multiple guide options for left and right guides. One each, for left and right, guides are chosen that will have high guide scores and least (or no) off target cleavage sites. **(b)** Schematic of donor DNA showing T7 promoter (as part of dsDNA template) and the actual part of ssDNA donor. Green arrow shows the primer for reverse transcription. **(c)** Floxed locus showing correct fusion of new sequence.

**Figure 4:**
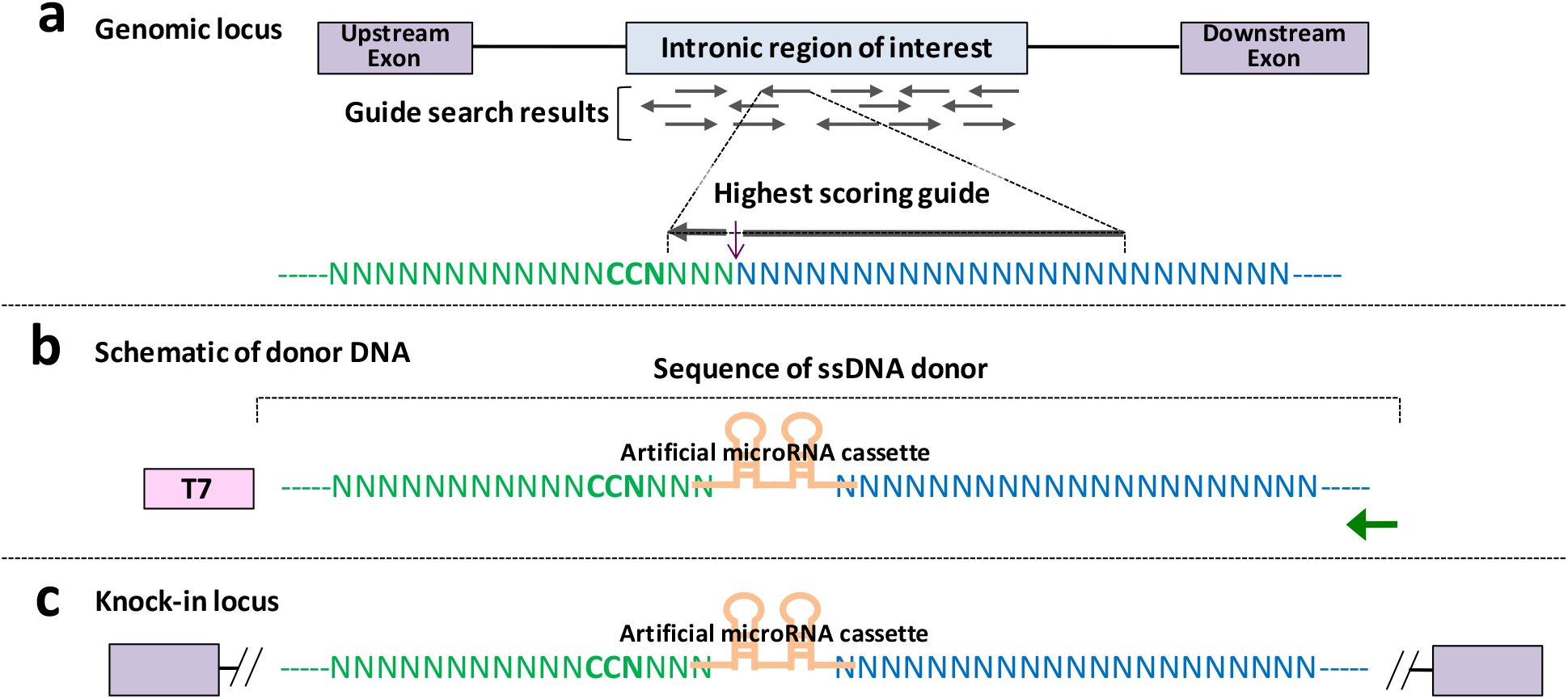
Design principles of inserting knock-down cassette using *Easi*-CRISPR and the architecture of ssDNA donor. **(a)** Genomic locus of a hypothetical gene’s target exon and its surrounding regions chosen for guide search. Hypothetical guide search results showing multiple guide options for left and right guides. One each, for left and right, guides are chosen that will have high guide scores and least (or no) off target cleavage sites **(b)** Schematic of donor DNA showing T7 promoter (as part of dsDNA template) and the actual part of ssDNA donor. Green arrow shows the primer for reverse transcription. **(d)** Knock-in locus showing correct fusion of new sequence.

#### The T7 promoter

The T7 prompter sequence is inserted immediately upstream to the left or right homology arms. The arm to which the T7 promoter is added is referred as upstream arm.

In our experiments thus far, we have successfully used either sense or anti-sense strands with respect to the direction of gene transcription.

#### The upstream homology arm

This corresponds to the upstream sequence from the point in the genome where the new sequence must be inserted (this is generally the sequence to the left of the cleavage point). We have not done a systematic study to identify optimal lengths for best performance; our initial studies contained arms from 55-105 bases long. It is preferable to choose a ‘G’ (ideally ‘GG’), at the 5’ end of the homology arm, to which the T7 prompter will be added upstream. It is known that presence of one or two Gs immediately 3’ to the T7 promoter increases T7 RNA polymerase transcription efficiency. **Note**: if a suitable sequence that matches this criterion is not available in the left homology arm, the T7 promoter can be added to the right homology arm, in which case the right homology arm will be referred to as the upstream homology arm (with respect to donor DNA direction).

#### The downstream homology arm

Corresponds to the downstream sequence from the point in the genome where the new sequence must be inserted (generally the sequence on the right of the cleavage point). Typically, about 55 to 105 bases long. A primer for reverse transcription is designed to bind at the 3’ end of downstream homology arm. Therefore, the terminal region should possess an optimal sequence for primer binding to be suitable for RT reaction (for example optimal GC content and containing non-repetitive sequences). A unique restriction site (that produces 5’ overhang or blunt end) should be added downstream of the right arm to linearize the plasmid before *in vitro* transcription.

#### The middle region

This region constitutes the new sequence to be inserted. The 3’ end of the upstream arm will continue to the 5’ end of the middle region (new sequence) and the 3’ end of the middle region will continue to the 5’ end of the downstream arm. For floxing designs (and for knock-down designs), once the theoretical sequences of the ssDNA cassette are designed, the cassettes can be built by custom synthesis from commercial vendors (such as Bio Basic, Integrated DNA Technologies, GENEWIZ, GeneArt, GenScript, or Life Technologies). For knock-in designs consisting of expression cassettes such as recombinases or reporters, any known plasmids can be used as templates to amplify the cassette using primers that contain the homology arms.

### Synthesis of ssDNA using *ivTRT* method

This method uses a dsDNA template to transcribe into RNA. The RNA is then reverse-transcribed back to DNA (to generate ssDNA molecules), followed by RNaseH degradation of RNA and purification of the ssDNA. The dsDNA templates can be either PCR products or plasmids that contain a T7 promoter and the insertion cassette (homology arms with the new sequence of interest in the middle). A schematic of ivTRT and ssDNA preparation steps is shown in **Figure 5**. Another method of generating ssDNAs was reported recently that uses DNA nicking endonucleases to make nicks on a plasmid dsDNA, followed by separation and purification of the desired ssDNA fragment from a denaturing agarose gel.^25^

**Figure 5:**
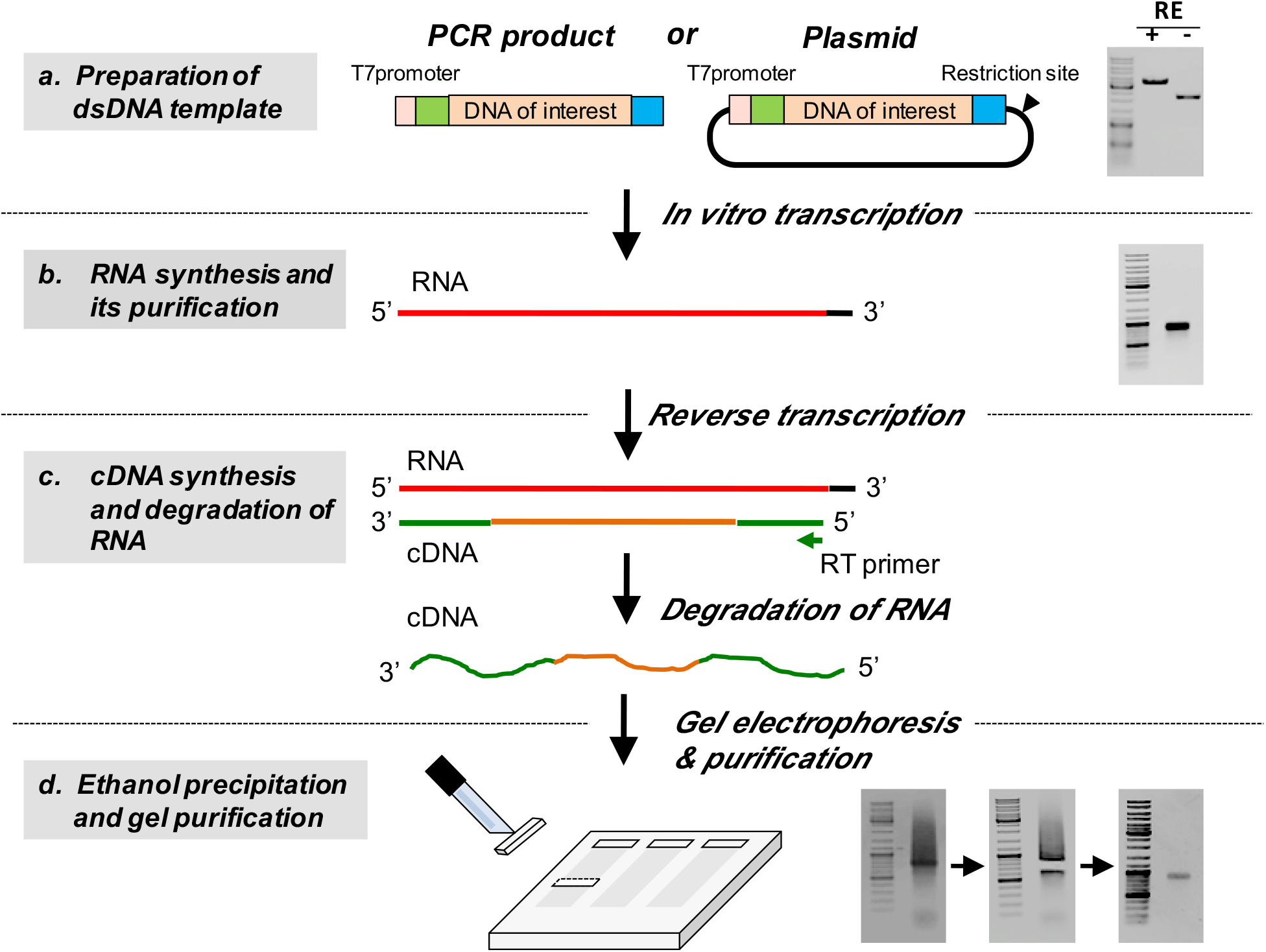
Schematic of *iv*TRT and ssDNA preparation steps. **(a)** A dsDNA template can be a PCR product or a plasmid with a suitable restriction enzyme (RE) site distal to the DNA cassette. The gel on the right shows a plasmid digested with a RE. (b) RNA is synthesized using *in vitro* transcription; the gel on the right shows an RNA of ~ 900 bases long. **(c)** cDNA is synthesized using a reverse primer. **(d)** Purification of ssDNA: The gel on the right shows running of ssDNA (cDNA) preparation. Note that cDNA preparation typically runs like a smear with a prominent band. The right half of the gel image shows the prominent band excised from the gel for purification. The lower gel image shows the purified ssDNA. The DNA size marker used in all the gels is GeneRuler DNA Ladder Mix (ThermoFisher Scientific, cat. no. SM0331).

### Preparation of injection mix

The ssDNA synthesized by ivTRT is then used as a repair donor for microinjection along with CRISPR components. Our first set of *Easi*-CRISPR method development experiments used Cas9 mRNA and single-guide RNA (sgRNA) as CRISPR components, but in our recent experiments we observed that RNAs that have been separated (crRNA and tracrRNA) and the Cas9 Protein (referred as ctRNP) have better rates of correct insertion. One reason for this could be that the pre-assembled ctRNP would be immediately available for cleavage upon injection, whereas there would be some delay in formation of Cas9 protein from the mRNA. A previous study showed that when comparing the sgRNA/Cas9 protein (sgRNP) and ctRNPs, ctRNPs had the highest knock-in efficiency.^26^ Another advantage of ctRNP composition is that all reagents can be commercially synthesized more cheaply than by preparing sgRNAs through *in vitro* transcription methods (Harms et al.; these were quite prevalent until about a year ago). To prepare *Easi*-CRISPR components for microinjection, ctRNP and ssDNA are mixed together; the process is described in detail in the “*Procedure*” section.

The microinjection step also involves a series of transgenic technologies that are typically performed at specialized core facility labs. These steps (involving numerous steps of CRISPR mouse genome engineering protocols), have been published many a times in the last two to three years,^23,24^ and have now become standard methods. We therefore omit these steps from the main protocol and have included them as a supplementary text file.

### Genotyping floxed and knock-in alleles

A general schematic of genotyping strategies is shown in **Figure 6**. At least three separate PCR genotyping reactions are suggested for Easi-CRISPR-derived offspring: one PCR each for the two *LoxP* insertions (in case of floxed alleles) and for the two junctions in case of knock-in and knock-down alleles, and the third PCR for the full-length PCR (in case of floxed alleles) and for insert-specific regions (in case of knock-in and knock-down alleles).

**Figure 6.**
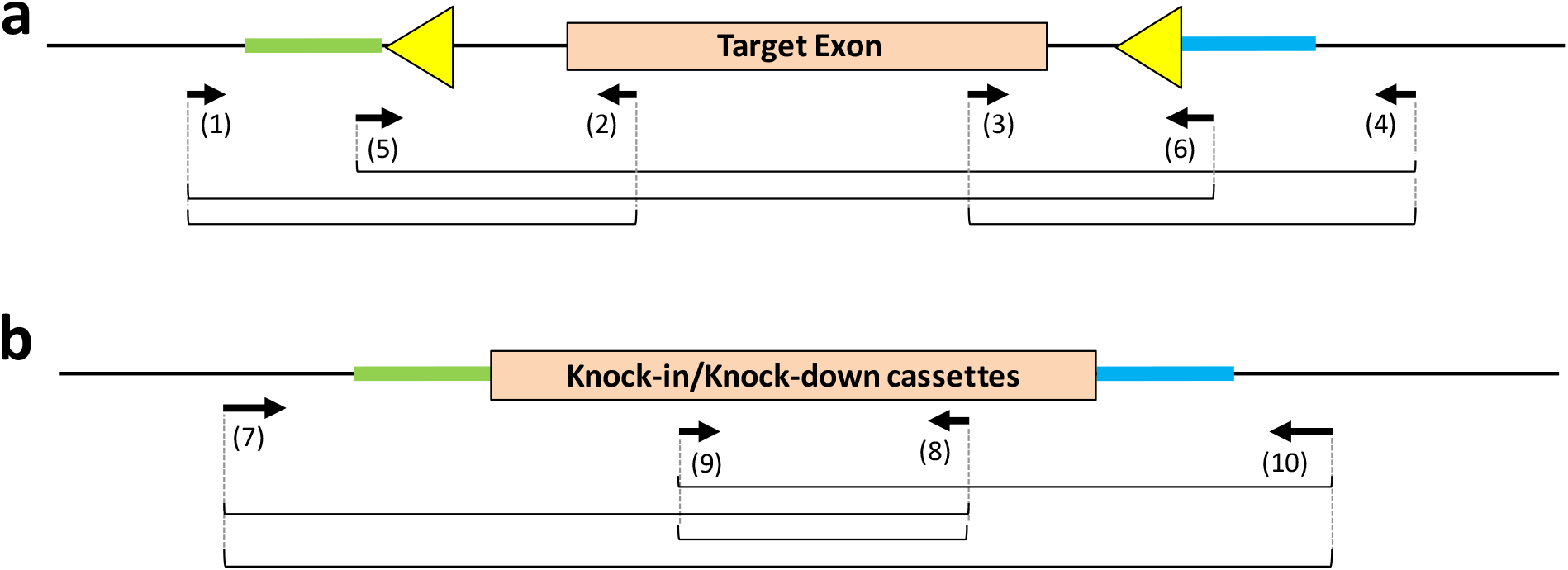
genotyping schematics. **(a)** Genotyping floxed alleles. Primer sets 1–2 and 3–4 amplify single *LoxP* insertion at the two separate sites but cannot indicate if they are inserted *in cis* or *in trans*. Correct insertion genotype (*in cis*) can be determined by PCR using primer sets (5–4 and 1–6). Note that 3’ regions of primers 5 and 6 bind to parts of *LoxP* sites (primer 5: 5’-NNNNNNNNNNNNNNNNNNNNataacttcgtatagc-3’ and primer 6: 5’-NNNNNNNNNNNNNNNNNNNNataacttcgtataat-3’). **(b)** Genotyping knock-in (and knock-down) alleles. One separate PCR each is performed for 5’ and 3’ junctional regions (primer sets 7-8 and 9-10) and for insert-specific regions (primer set 9-8). PCR with outer primer set (7-10) amplifies longer PCR fragments, including the entire knock-in cassette, as well as a shorter product originating from non-inserted wild type (or *indel*) allele. Positions of primers 9 and 8 (within the cassette) can be switched to reduce the amplicon sizes of junctional PCRs, in which case another set of insert specific primers need to be used in place of 9 and 8). The amplified fragments in both (a) and (b) should be sequenced to ensure sequence fidelity.

#### Floxed allele genotyping

The primers that can amplify each of the two separate *LoxP* sites will be useful for detection. When genotyping floxed alleles, we always use primer sets that amplify single *LoxP* insertion (for both *LoxPs*). Because the LoxP-inserted allele can be seen as a slightly larger band compared to the wild type, PCR products from wild type control should be included in agarose gel electrophoresis. When both *LoxPs* are positive, these pups can be candidates for floxed mice. However, it is often difficult to judge whether these two *LoxPs* are inserted *in cis*. Complete insertion of a floxed cassette *(in cis)* can be determined by PCR using primer sets (one outer primer and one other primer that bind to a distantly located *LoxP* site), and following sequencing of the amplified fragments to determine whether the fragment contains an internal *LoxP* site. Even when the correctly inserted floxed cassette is identified, it is desirable to check for the presence of a floxed allele and its nucleotide sequence in the next generation. The recombination capability of the floxed allele is confirmed by breeding the mouse line with a Cre driver line. A quick analysis for recombination of the floxed allele can also performed using an *in vitro* Cre reaction, where the target region is PCR-amplified and incubated with a Cre protein, and the recombination products are analyzed on agarose gel,^6^

#### Knock-in and knock-down alleles genotyping

We recommend PCR be performed for both the 5’ and 3’ junctional regions and for insert-specific regions (using internal primer sets). Even though all three sets of PCR reactions can show expected amplicons, all junctions and the insertion cassette must be fully sequenced to ensure sequence fidelity, to rule out any *indels* near the junctions, and to check for nucleotide mis-incorporations in the insert regions that may arise from the enzymatic synthesis steps in preparing the ssDNA donor. PCR with an outer primer set (that bind outside the homology arms) amplifies longer PCR fragments, including the entire knock-in cassette, as well as the shorter wild-type sequence (or not inserted but with *indel* derived from NHEJ). In this PCR, amplification of only the wild-type (or *indel*) band does not always guarantee that it can serve as a negative sample, because a smaller amplicon (wild-type) typically competes amplification of the longer band (from the correctly targeted allele). The insertion allele (larger band) may be amplified efficiently if the insertion event is biallelic (with no wild type allele). Regardless of the PCR amplification results, however, sequencing of the entire cassette is mandatory to ensure the correctness of the targeted allele. Sequencing should also be performed in F1 pups to rule out any hidden mosaic alleles that may contain mutations from the founders and transmitted to the offspring.

### Limitations of *Easi*-CRISPR

The Easi-CRISPR strategy allows for generation of various kinds of mouse models at an efficiency much higher than that of any previously described methods. Some of the limitations of Easi-CRISPR method are discussed below:

#### The length of the ssDNA donor

One of the current technical limitations of Easi-CRISPR is the synthesis of longer ssDNA molecules. We have successfully made ssDNAs up to ~2kb, which is sufficient for most of the commonly needed mouse models. Because of the enzymatic amplification steps involved in ivTRT method, some difficult-to-amplify type of sequences may be challenging to synthesize.

#### Possible sequence mis-incorporations arising during ssDNA preparation steps

Because of enzymatic amplifications steps, nucleotide mis-incorporations can occur during ssDNA synthesis. Some of the ssDNA preps at the beginning stage of developing this method had such mutations. We presume that a portion of these mutations may have resulted from the unintended use of standard grade enzymes rather than high-fidelity enzymes. Some knock-in mouse models generated may contain such incorporations, and therefore, the full cassettes must be sequenced among the founders and the germ line transmitted F1 offspring (to exclude the possibility of undetected mutations transmitted due to mosaicism). Although most reverse transcriptase enzymes do not have proofreading capabilities, the development of enzymes with proofreading function will be helpful to prevent mis-incorporations^27^.

#### Possible random insertion of ssDNA donors

Even though ssDNAs are less likely to be inserted into the genome at random locations, the possibility of random insertions cannot be excluded completely. Very rarely, some negative founders (as judged by the two junction PCR results) showed positive amplicons using internal primers suggestive of random insertion events.

## MATERIALS

### REAGENTS

#### CRISPR REAGENTS

- crRNA (med-mod) and tracrRNA (cat. no. 1072534) from Integrated DNA Technologies (IDT). The crRNA part is custom synthesized for each specific guide RNA, whereas tracrRNA is universal
- Alt-R^™^ S.p. Cas9 Nuclease 3NLS (cat. no. 1074181) from IDT

#### REAGENTS FOR ssDNA SYNTHESIS

General note: listed below are the standard kits and reagents that have been tried in our laboratories. Comparable kits and reagents from other vendors may also be used in place of these.

- Standard desalt Ultramers (long primers) from IDT (or any commercial vendor) to use as primers for PCR of insert or custom gene-synthesized plasmid (any commercial vendor)
- KOD plus neo (TOYOBO, cat. no. KOD-401) or any suitable high-fidelity PCR mixes
- dNTP mix (New England Biolabs, cat. no. N0447L)
- SeaKem ME Agarose (Lonza, cat. no. 50011) and SeaPlaque GTG Agarose (Lonza, cat. no. 50110)
- Wizard SV Gel PCR Clean up System kit (Promega, cat. no. A9282) or NucleoSpin Gel and PCR Clean-up (MACHEREY-NAGEL, cat. no. 740609)
- MEGAclear Kit (Ambion-Life Technologies. cat. no. AM1908)
- T7 RiboMax Express Large Scale RNA Production System (Promega, cat. no. P1320)
- Ethanol 200 Proof ACS Grade (Deacon Laboratories) or Ethanol (99.5) (Wako, cat. no. 057-00456)
- M-MuLV Reverse Transcriptase (New England Biolabs, cat. no. M0253S) or SuperScript III Reverse Transcriptase (Life Technologies, cat. no. 18080051) or SuperScript IV Reverse Transcriptase (Life Technologies, cat. no. 18090010)
- RNaseH (New England Biolabs, cat. no. M0253S)
- S1 nuclease (TaKaRa, cat. no. 2410A)
- PCR tubes (ThermoFisher Scientific, cat. no. AB-1114)
- RNase free Pipet Tips (ART-Aerosol Resistant Tips)
- 3mol/l-Sodium Acetate Buffer Solution (pH5.2) (Nakalai Tesque, cat. no. 06893-24)
- Phenol, Saturated with TE buffer (Nakalai Tesque, cat. no. 26829-54)
- Modified TE: 10mM Tris-HCl, 0.1mM EDTA, pH8.0 (e.g., Affymetrix, cat. no. 75793)
- Embryo Max Microinjection Buffer (EMD Millipore, cat.no. MR-095-10F)
- MILLEX-GX 0.22μm Filter Unit (EMD Millipore, cat.no. SLGV004SL)

#### REAGENTS FOR MOUSE TRANSGENSIS EXPERIMENTS

These experiments follow well-established, standard protocols of mouse transgenesis, typically performed at specialized core facility laboratories. Such protocols have been described in detail elsewhere (Refs). Details of these reagents and procedures are included in the supplemental text.

#### REAGENTS FOR MOUSE GENOTYPING

- Cell Lysis Solution (Qiagen, cat. no. 158908)
- Protein Precipitate Solution (Qiagen, cat. no. 158912)
- DNA Hydration Solution (Qiagen, cat. no. 158914)
- Go Taq Hot Start Green Mix for genotyping (Promega, cat. no. M5122-23)
- Allele-In-One Mouse Tail Direct Lysis Buffer (KURABO, ABP-PP-MT01500)
- Agarose (Phenix Research Products, RBA 500)
- 50X TAE Buffer (ThermoFisher Scientific, cat. no. BP1332-20)
- Water DEPC-treated Nuclease free (ThermoFisher Scientific, cat. no. BP561-1)
- TA Cloning^®^ Kit, with pCR™2.1 Vector (Life Technologies, cat. no. K202020)
- LB agar Ampicillin-100, Plates (Sigma, cat.no. L5667-10EA)
- Mix & Go Competent cells – Strain Zymo 5α (Zymo Research, cat.no. T3007)
- QIAprep Spin Miniprep Kit (Qiagen, cat. no. 27106)
- Proteinase K solution (10ml; 5Prime, cat. no. 2500150)
- GeneRuler DNA Ladder Mix (ThermoFisher Scientific, cat. no. SM0331)
- Gel Loading Dye Purple (6X), no SDS (New England Biolabs, cat. no. B7025S)
- Ethidium bromide (Sigma, cat. no. E1510-10ml)
- Kimwipes for general cleaning (Kimtech Science, cat. no. KCK-280-4.4 × 8.4)

### EQUIPMENT

- Thermocycler (BioRad T100 or equivalent)
- UV spectrophotometer (NanoDrop-1000, Thermo Scientific) Gel documentation System (Gel Doc XR+, BioRad)
- LED light (BioSpeed ethidium bromide-VIEWER)
- Tabletop micro-centrifuges (Model Eppendorf 5417C or equivalent)
- Tabletop micro-centrifuges-refrigerated (Model Eppendorf 5417R or equivalent)
- Micropipettes (Eppendorf Research Plus)
- Gel Electrophoresis and EPS (Electrophoresis Power Supplies or equivalent)
- Shaking heat block (Eppendorf, Thermomixer Comfort)
- Heat block (TAITEC, Dry ThermoUnit)
- Water baths (Fisher Scientific)
- Vortex Genie 2 (Scientific Industries, Inc)
- Microwave and weighing balances

### REAGENT SETUP

#### Oligonucleotides and Ultramers

- Re-suspend oligonucleotides or ultramers to a final concentration 100μM in nuclease free water and can be stored at −20°C for future use.

#### 1X TAE Electrophoresis Buffer

- Dilute 50XTAE buffer in distilled water to a 1X working solution. 1X solution can be stored at room temperature for up to 3 months.

#### Ethanol

- For 100ml of ethanol (70% (vol/vol)) solution, combine 70 ml 99.5% (vol/vol) ethanol with 30 ml of Nuclease free water and store it in a tightly sealed tube at room temperature.

#### crRNA and tracrRNA

- Re-suspend crRNA and tracrRNA to a final concentration 100μM (approx. 1.2μg/μl for crRNA and 2.2μg/μl for tracrRNA) in Injection buffer and can be stored at −80°C for future use.

#### The ctRNP complex preparation

- CRISPR system comprises three components: crRNA, tracrRNA, and Cas9 Protein. The crRNA is unique to each project, whereas the other two components are universal. The crRNA is 36 bases long and tracrRNA is 67 bases long. The crRNA can be custom synthesized (e.g., Alt-R system from IDT); they can also be chemically modified, which is known to increase their stability^28^. We have used minimum and medium modifications of crRNAs thus far, with no apparent differences in editing efficiencies. The two RNAs (crRNA and tracrRNA) are annealed at 1:1 molar concentration to generate active guide RNA and the guideRNA is mixed with Cas9 protein to obtain ctRNP complexes.

#### Cas9 Protein

- Cas9 protein is diluted to (e.g. 3.1μM [500ng/μl]) with injection buffer and aliquots are stored at −20°C for future use.

#### Preparation of MILLEX-GX (0.22μm Filter Unit) for steps 29–35

- Cut a PCR tube from an 8 strip PCR tube and place the filter unit MILLEX-GX (0.22μm Filter Unit) into the PCR tube (**Fig. 7**). Insert the column (filter unit in the PCR tube) into 1.5ml micro-centrifuge tube.

**Figure 7.**
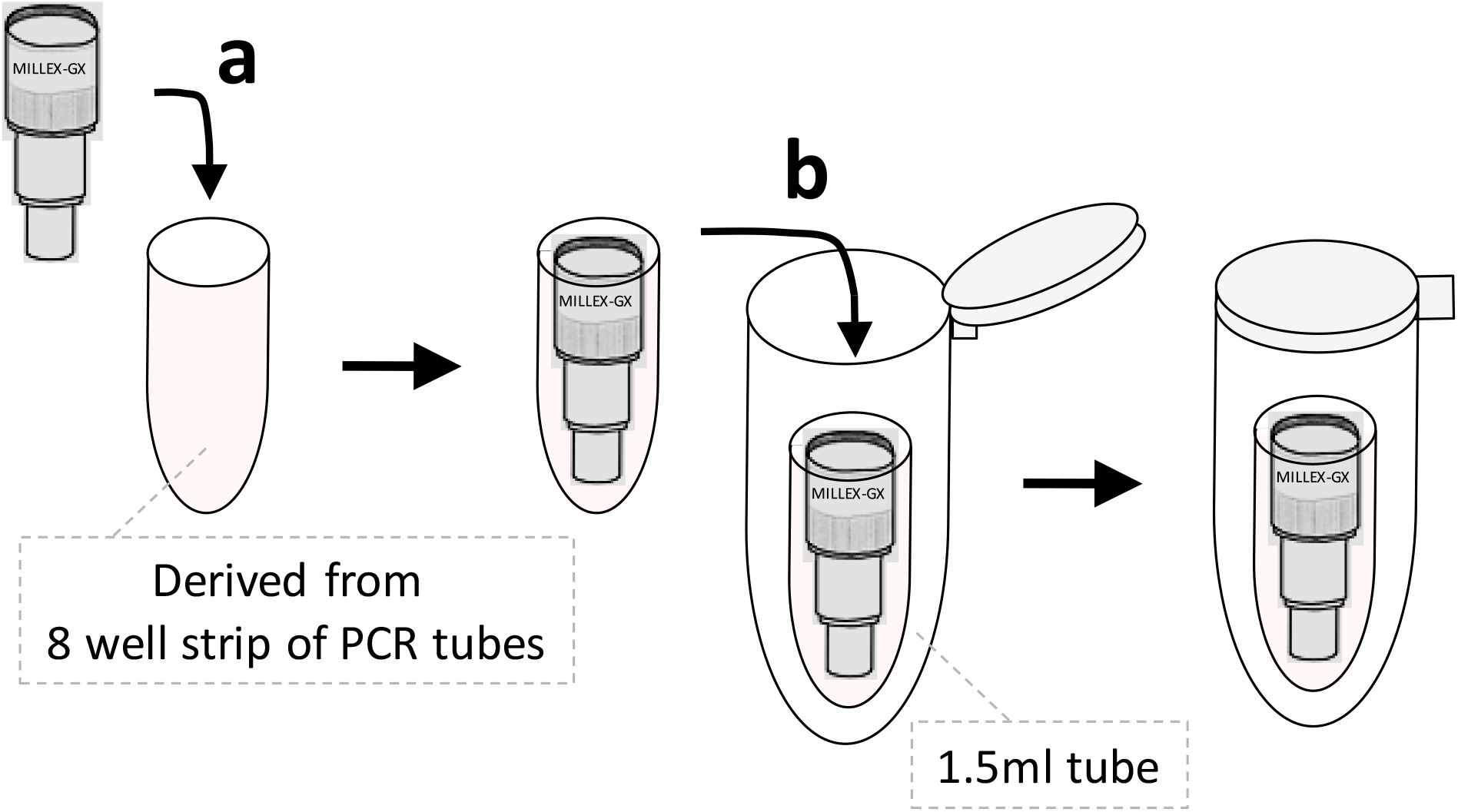
Assembling of MILLEX-GX 0.22 μM filter unit. **(a)** insert a Millex-Gx 0.22 μM filter unit into a PCR tube (cut from an 8 well strip) **(b)** Insert the PCR tube containing the filter into a 1.5ml micro-centrifuge tube.

## PROCEDURES

### Steps 1 – 2 Preparation of dsDNA template ● TIMING 5h

**1)** If using plasmid DNA as templates, digest appropriate amount of 2.5–4μg of plasmid DNA (see options A and B below) with restriction enzyme (RE) for linearization. Optional: If using PCR products as templates (for RNA synthesis), generate the product using high fidelity Taq DNA polymerase. Primers used for amplifying PCR products can be standard desalt grade from any commercial vendor.

Option A: if linearized plasmid is used: Use about 2.5-4μg of plasmid for digestion in a volume of 100μl.
Option B: if PCR product is used purified using gel purification: Use about 5–15μg of PCR product to load the gel **▲ CRITICAL STEP** Use Hi-Fidelity Taq polymerase for generating template using PCR. After RE digestion of the plasmid or generating the PCR products, verify the samples for digestion or PCR amplification by agarose gel electrophoresis. Load ~1–2μl of RE digested sample or PCR Product using 1% (SeaKem ME agarose gel) in 1× TAE.
**2)** Purify the dsDNA template prepared in step 1:

a. Option A: if linearized plasmid is used as a template:

i. Perform phenol extraction and ethanol precipitation and finally re-suspend it in 2μl of modified TE buffer.
b. Option B: if PCR product is used as a template:

i. Gel purify the PCR product using (Wizard SV Gel PCR Clean up System kit) and perform two elutions in 20μl Elution buffer. Estimate the concentration using Nano Drop. **! CAUTION** Phenol is toxic and cause burns. Should be opened in fume hood wearing proper protective equipment.

### Steps 3 – 5 RNA synthesis using T7 RiboMax Express ● TIMING 1-4h

**3)** RNA is synthesized according to manufacturer’s instructions (T7 RiboMAX Express Large Scale RNA Production System). Add the following reagents into a PCR tube or micro centrifuge tube and mix by pipetting and give a short spin. About 1–2μg of template DNA is used for *in vitro* transcription. **! CAUTION** Avoid repeated freeze-thaw of buffers. Thaw all the reagents on ice and be sure to use RNase-free tips and gloves.

**4)** Incubate the reaction at 37°C in a thermocycler for 30 minutes to 3 hours.
**5)** Add 1μl of RQ1 RNase-Free DNase, mix well, and incubate at 37°C in a thermocycler for 15 minutes to eliminate the DNA template. **? TROUBLESHOOTING** **! CAUTION** Wrap tube caps with parafilm to prevent contamination from water bath.

| Component | Amount per Reaction (µl) |
| --- | --- |
| RiboMAX Express T7 2×Buffer | 10µl |
| Template DNA | ..µl* |
| Nuclease free water | ..µl |
| Enzyme Mix, T7 Express | 2µl |
| Total | 20µl |

\* If template from Step 2a (Option A) is used, use the 1 µl. If template from Step 2b (Option B) is used, adjust the µls according to the DNA concentration of the eluate

### Steps 6–18 Purification of RNA using MEGAclear kit ● TIMING 1h

**6)** RNA is purified using MEGAclear Kit according to manufacturer’s instruction.
**7)** Preheat a dry heat block at 65–70°C and pre-warm the elution solution.
**8)** Add 80μl of elution solution to the sample (total volume to 100μl) and mix by gentle pipetting.
**9)** Add 350μl of Binding Solution and mix by gentle pipetting.
**10)** Add 250μl of >99.5% ethanol and mix by gentle pipetting.
**11)** Transfer the sample to the column and centrifuge (13,500–14,000rpm,1 minute, room temperature).
**12)** Discard the flow-through and re-insert the column into the micro-centrifuge tube.
**13)** Add 500μl of washing solution to the column and centrifuge (13,500–14,000rpm, 1 minute, room temperature).
**14)** Discard the flow-through and repeat washing (as in step 13).
**15)** Discard the flow-through and centrifuge (13,500–14,000rpm, 1 minute at room temperature) to completely remove traces of ethanol.
**16)** Insert the column into newly labelled 1.5ml micro-centrifuge and add 25–50μl of prewarmed elution solution directly into the column bed, incubate for 10 minutes, and Centrifuge at 13,500–14,000rpm, 1 minute, at room temperature). A second elution can be performed to recover more RNA.
**17)** Check the quality and concentration of RNA by analyzing 1μl of sample in a Nano Drop.
**18)** Confirm the quality of RNA by agarose gel electrophoresis (**Fig. 5b**). **■ PAUSE POINT** Store the samples at −80°C until use. **▲ CRITICAL STEP** Aliquot about 3–5μg RNA/tube to avoid repeated freezing and thawing of RNA. **? TROUBLESHOOTING**

### Steps 19–23 Synthesis of cDNA from RNA ● TIMING 1.5h

**19)** cDNA is synthesized according to manufacturer’s instructions (Superscript III RT protocol). Add the following reagents to a PCR tube or micro-centrifuge tube (Tube A) and mix by pipetting and give a short spin. About 3–5μg of purified RNA is used as template for cDNA Synthesis.

**20)** Incubate the tube at 65°C for 5minutes.
**21)** Immediately place the tube (Tube A) on ice, wait for at least one minute and then proceed to cDNA synthesis. Mix the following reagents in a new tube (Tube B).

| Component | Amount per Reaction (µl) | Final (in 30µl) |
| --- | --- | --- |
| RNA Template | ..µl | ~5µg |
| RT Primer (100µM) | 1.5µl | 5µM |
| dNTP Mix, NEB (10mM) | 3.0µl | 1mM |
| Nuclease free water | ..µl |  |
| Total | 30µl* |  |

\* Reactions can be scaled up to obtain higher quantity of cDNA. This reaction is 3x scale (3 X 10 µl) of manufacturer's protocol. Mix the reaction by pipetting and short spin in centrifuge.

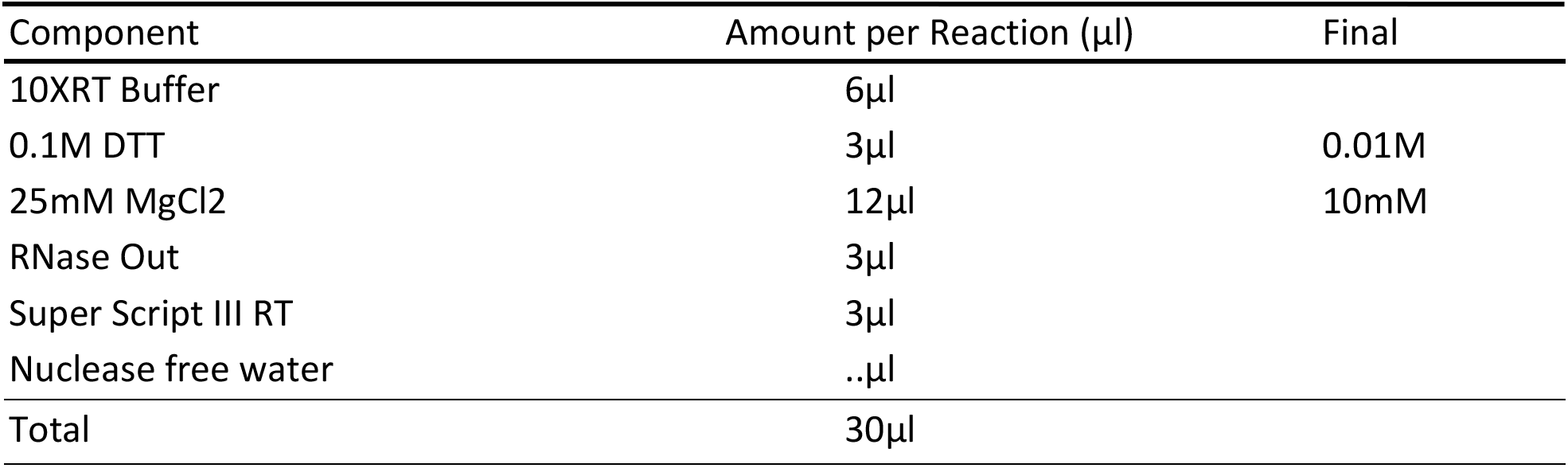 **! CAUTION** DTT is harmful when inhaled, it may cause irritation to skin and eye. Wear Gloves and proper equipment when working with DTT.
**22)** Combine contents of both tubes (Tubes A and B). Mix well and incubate the reaction at 50°C for 50 minutes in a water bath or thermocycler. Stop the reaction by heat-inactivating at 85°C for 5 minutes. Cool down the reaction to room temperature.
**23)** Add 3μl of RNaseH to the tube, mix by pipetting and incubate it at 37°C for 20 minutes in water bath. **? TROUBLESHOOTING** **! CAUTION** Wrap the tube caps with parafilm to prevent any contamination from water bath. ■ **PAUSE POINT** The samples can be stored at −20°C until use.

### Steps 24 – 28 Ethanol precipitation of cDNA and gel purification ● TIMING 1.5h

**24)** Add 40μl of modified TE (10mM Tris-HCl, 0.1mM EDTA), 10μl of 3M Sodium acetate (pH5.2) and 250μl of 99.5% ethanol to 60μl of sample. Vortex and centrifuge at 14,000rpm, 4°C, 10 minutes. Discard the supernatant, add ~300μl 70% ethanol and centrifuge (14,000rpm, 4°C, 2 minutes).
**25)** Completely remove the supernatant and air-dry the pellet. Dissolve the pellet in 8μl of modified TE.
**26)** Add 2μl of 6x gel loading buffer, load the sample in to 1% low-melting gel (SeaPlaque GTG agarose), and perform electrophoresis (135V, 30 minutes).
**27)** Stain the gel with ethidium bromide, excise the gel piece containing the major band under LED light (BioSpeed ethidium bromide-VIEWER), and transfer it to a new 1.5ml microcentrifuge tube. A sample electrophoresis image of a cDNA preparation and excising of a prominent band for gel purification is show in **Figure 5c**. **! CAUTION** (i) Ethidium Bromide is carcinogenic and mutagenic. Wear personal protective equipment to avoid exposure to UV light. (ii) Suggest using long-wave length UV to prevent damage to the DNA sample. The DNA from the gel slice can be extracted using one of the below two options:

a. Option A: Using NucleoSpin Gel and PCR Clean-up (TaKaRa) kit.
b. Option B: Phenol extraction and ethanol precipitation.
**<u>28) Option A: Using NucleoSpin Gel and PCR Clean-up (Column purification)</u>**

- NucleoSpin Gel and PCR Clean-up (TaKaRa) used for purification of cDNA according to manufacturer’s instruction.
- Add 200μl Buffer NT1 to each 100mg of agarose gel (containing cDNA) and mix well. Determine the weight of the gel piece before adding Buffer NT1.
- Incubate the tube for 10 minutes at 50^o^C to completely elute gel piece (briefly vortex the sample every 2–3 minutes).
- Transfer the sample (up to 700μl) into a column and centrifuge for 30 seconds at 10,000rpm.
- Discard flow-through and add 700μl Buffer NT3 to the column and centrifuge for 30 seconds at 10,000rpm.
- Repeat washing with Buffer NT3 (above step).
- Centrifuge for 1 minute at 10,000rpm to completely remove Buffer NT3.
- Place the column into a new micro-centrifuge tube, add 15–30μl Buffer NE, and incubate at room temperature for 1 minute.
- Centrifuge for 1minute at 10,000rpm [1st elution].
- Place the column into the tube and add 15–30μl Buffer NE and incubate room temperature for 1 minute.
- Centrifuge for 1 minute at 10,000rpm [2nd elution] (total elution volume will be 30–60μl). ▲ **CRITICAL STEP** Multiple elution yields higher cDNA recovery; for example, three elutions of 20 μl each can also be performed.
**<u>28) Option B: Phenol extraction and ethanol precipitation</u>**

- Add 200μl of modified TE into the gel piece-containing tube and place it at −80°C for more than 20minutes (can be left overnight).
- Melt the sample at room temperature and transfer the solution to a new tube.
- Add 200μl of TE-saturated phenol and centrifuge (12,000rpm, 20°C, 6 minutes). **! CAUTION** Phenol is toxic and cause burns. Should be opened in fume hood wearing proper protective equipment.
- Transfer the supernatant to a new tube, add 20μl of 3M sodium acetate (pH5.2) and 500μl of 99.5% ethanol, vortex and centrifuge at 14,000rpm, 10minutes, at room temperature.
- Decant supernatant, add 130μl of 70% ethanol, centrifuge at 14,000rpm, 2minutes, 4°C.
- Completely remove the supernatant and dry the pellet. Dissolve the pellet in 11μl of injection buffer. ■ **PAUSE POINT** Store the samples at −20°C (short term) or −80°C (long term). **? TROUBLESHOOTING**

### Steps 29–35 Filter purification of cDNA solution

**29)** Pre-cool the centrifuge at 4°C.
**30)** Add injection buffer to the MILLEX-GX (0.22μm Filter Unit: see reagent setup for assembling a filter) and centrifuge the tube (11,200rpm, 4°C, 1 minutes).
**31)** Take the filter unit only from the tube and place it in a new PCR tube, re-insert to 1.5ml micro-centrifuge tube.
**32)** Add the 11μl of sample on the filter and centrifuge it (11,200rpm, 4°C, 2 minutes).
**33)** Take the filter-containing PCR tube from the 1.5ml micro-centrifuge tube and transfer the filtered sample (in a bottom of the PCR tube) solution to a new 1.5ml tube.
**34)** Transfer 1μl of sample to another new tube and dilute with 4μl injection buffer (total 5μl). Use 1μl for concentration determination using Nano Drop, and remaining 4μl for agarose gel electrophoresis (**Fig. 5d**).
**35)** Store the remaining sample (10μl) at −20°C or −80°C for future use. **? TROUBLESHOOTING**

### Step 36 Preparation of guide RNA by annealing crRNA and tracrRNA ● TIMING 30 minutes

**36)** Anneal crRNA and tracrRNAs by mixing equi-molar ratios. Mix 10μl of 6.1μM crRNA (approx. 72ng/μl) and 10μl of 6.1μM tracrRNA (approx. 135ng/μl) and anneal in a thermocycler (94°C for 2 minutes and then place at room temperature, about 10 minutes). The crRNA/tracrRNA complex is then stored at −80°C until use. **? TROUBLESHOOTING** ▲ **CRITICAL STEP** Annealing of crRNA and tracrRNA is an important step to obtain guide RNA and for the formation of ctRNP complex. Since the Cas9 protein is supplied at high concentrations, we occasionally make intermediary dilutions (e.g. 3.1μM [500ng/μl]) with injection buffer and then dilute it to final concentration (0.3μM [50ng/μl]), as described below.

### Steps 37–41 Preparation of ctRNP + ssDNA Injection mix ● TIMING 1h

**37)** Preparation of ctRNP injection mix (10–20μl volume). Cas9 protein is diluted with injection buffer to the concentration of 500 ng/μl (equivalent to 3.1μM).
**38)** Mix the components on ice as below (10μl scale).

**39)** We use 0.6μM (20.7ng/μl) of guide RNA and 0.3μM (50ng/μl) of Cas9 protein for mouse zygote microinjections. Other concentrations may also work. **? TROUBLESHOOTING**
**40)** Incubate injection mixture at 37°C for 10 minutes just before injection and keep it at room temperature until use.
**41)** Load injection mix into needles and follow microinjection procedures described in Harms et al. (2014) [1], References 1. Harms DW, Quadros RM, et al. (2014) Mouse Genome Editing using CRISPR/Cas System. Current protocols in human genetics/editorial board, Jonathan L. Haines [et al.]. 83:15.17.11–15.17.27.

| Component | Amount per Reaction (µl) | final concentration |
| --- | --- | --- |
| crRNA/tracrRNA complex (3.1µM [104ng/µl]) | 2µl | 0.6µM (20.7ng/µl) |
| ssDNA (30-100ng/µl) | ..µl | 5-10ng/µl |
| Cas9 protein (3.1µM [500ng/µl]) | 1µl | 0.3µM (50ng/µl) |
| injection buffer | ..µl |  |
| Total | 10µl |  |

### Steps 42–43 Genotyping of offspring to identify transgenic founders

**42) <u>Mouse tail or ear DNA extraction</u> ● TIMING 1d** Described below are two methods for mouse DNA extraction: the longer protocol produces cleaner DNA, and the rapid protocol produces crude DNA but can be used for quick genotyping. DNA preparations from both methods work well for genotyping PCRs. **<u>Longer protocol:</u>**

i. Collect ~2–3mm tail pieces from mice in 1.5ml micro-centrifuge tubes. Collect a sample from a wild type mouse as control. Add 300μl Cell Lysis Solution containing 3μl Proteinase K and incubate at 65°C overnight in a dry bath. **▲ CRITICAL STEP** To save time and reagent loss, make a master mix for the lysis solution and Proteinase K. This can then be distributed to 300μl aliquots in separate micro-centrifuge tubes.
ii. Next day cool the samples at room temperature. Add 100μl of the Protein Precipitation Solution and vortex thoroughly for ~20 seconds.
iii. Place the tubes on at 4°C for 10 minutes, and then centrifuge at 13,000 rpm for 5 minutes.
iv. Transfer supernatants to newly labeled micro-centrifuge tubes containing 800μl of 100% ethanol. Mix by inverting the tubes 8–10 times. **▲ CRITICAL STEP** Use ethanol resistant markers for labeling the tubes because spills of ethanol solutions on the tubes can wipe away the labels.
v. Centrifuge at 13,000 rpm for 5 minutes.
vi. Slowly discard the supernatant, add 700 μl of 70% ethanol, mix by inverting 8–10 times.
vii. Centrifuge at 13,000 rpm for 5 minutes, then slowly discard supernatant.
viii. Centrifuge at 13,000 rpm for 4 minutes.
ix. Aspirate remaining 70% ethanol using a 200μl pipette tip and air dry DNA pellet for ~5 minutes (do not exceed minutes). **? TROUBLESHOOTING** **! CAUTION** It is important to change tips between samples to avoid cross-contamination. **▲ CRITICAL STEP** The DNA pellet is very loose at this step; the aspiration step should be performed with care to avoid losing the pellet.
x. Add 100μl DNA Hydration Solution to the pellet and flick the side of the tube to mix. Incubate tubes at 65°C in dry bath for 15–30 minutes to solubilize the DNA. **■ PAUSE POINT** Genomic DNA samples can be stored at 4°C for later use. **Rapid protocol:**

i. Collect ~2mm ear pieces from mice in 1.5ml micro-centrifuge tubes. Collect a sample from a wild type mouse as control. Add 40μl Allele-In-One Mouse Tail Direct Lysis Buffer and incubate at 55°C with shaking using shaking heat block more than 3 hours, or overnight.
ii. Incubate tubes at 85°C in a heat block for 45 minutes to inactivate the protease in the solution. **■ PAUSE POINT** Genomic DNA samples can be stored at −20°C.
iii. in Centrifuge at 14,000rpm for 2 minutes (because the samples are crude) and use for PCR reactions.
**43) <u>PCR amplification and agarose gel electrophoresis ● TIMING 1d</u>**

i. Perform PCR for all primer sets on all samples including a DNA sample from wild type mouse as a control.
ii. Use the 2X PCR master mix manufacturer’s protocol to determine optimal parameters for your PCRs. The basic steps for GoTaq Hot Start Green Master Mix are outlined below:
iii. Combine PCR master mix, nuclease-free water, mouse tail DNA (~1μl of Template, from genomic DNA Isolation), and primer mix (both forward and reverse) for a total volume of 20μl/sample.
iv. Run the PCR reactions in a thermocycler; the following are standard PCR conditions:

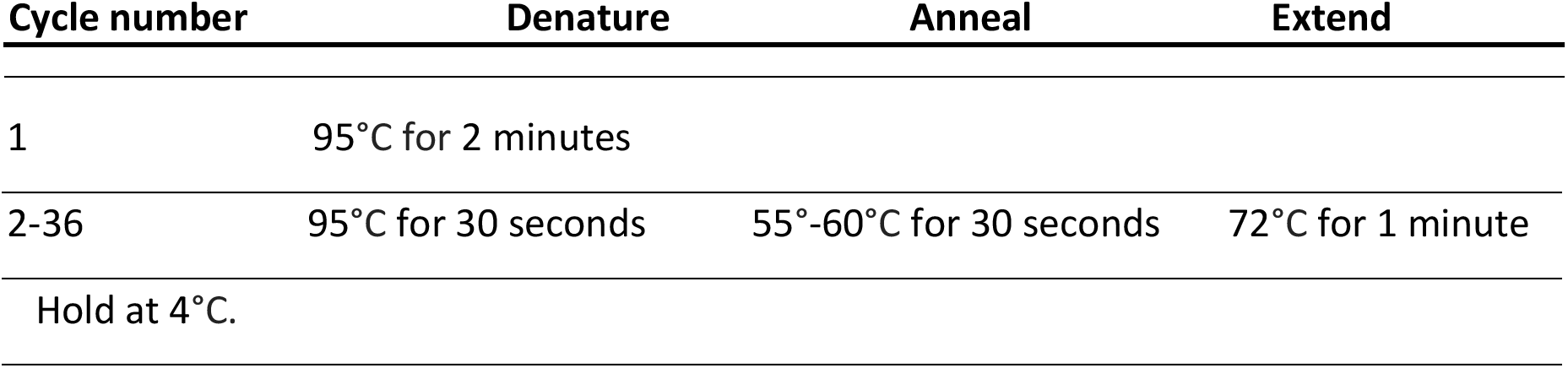
v. Run PCR products on a 1% agarose gel. Analyze gels to interpret genotyping results.
vi. Sequence PCR products directly, or clone PCR products by TA Cloning kit and sequence them to confirm the genotypes. **? TROUBLESHOOTING**

| Cycle number | Denature | Anneal | Extend |
| --- | --- | --- | --- |
| 1 | 95°C for 2 minutes |  |  |
| 2-36 | 95°C for 30 seconds | 55°-60°C for 30 seconds | 72°C for 1 minute |
| Hold at 4°C. |  |  |  |

## TIMING

A typical *Easi*-CRISPR mouse genome engineering project can be completed in about 2 months from the time the CRISPR strategy is designed up to identifying the genome-edited G0 founder pups. The general time frame required for different stages of *Easi*-CRISPR are outlined in **Figure 1**. Even though designing the overall strategy (first stage) can only take about 2–3 hours for an experienced technician (twice as much or more for others), this is the most critical part of the *Easi*-CRISPR protocol. Even slight errors in designing the donor, choosing the guides, and other steps at this stage can result in a mouse model that may not be developed exactly as intended. We recommend that two or three independent persons look at the design, and that at least one of these be an expert with previous successful experience in designing genetically engineered models. The next critical step is the preparation of ssDNA donors, which requires the most hands-on time of the *Easi*-CISPR protocol steps. Timing considerations for these steps are described, including optional pause points in the protocol steps.

### Troubleshooting

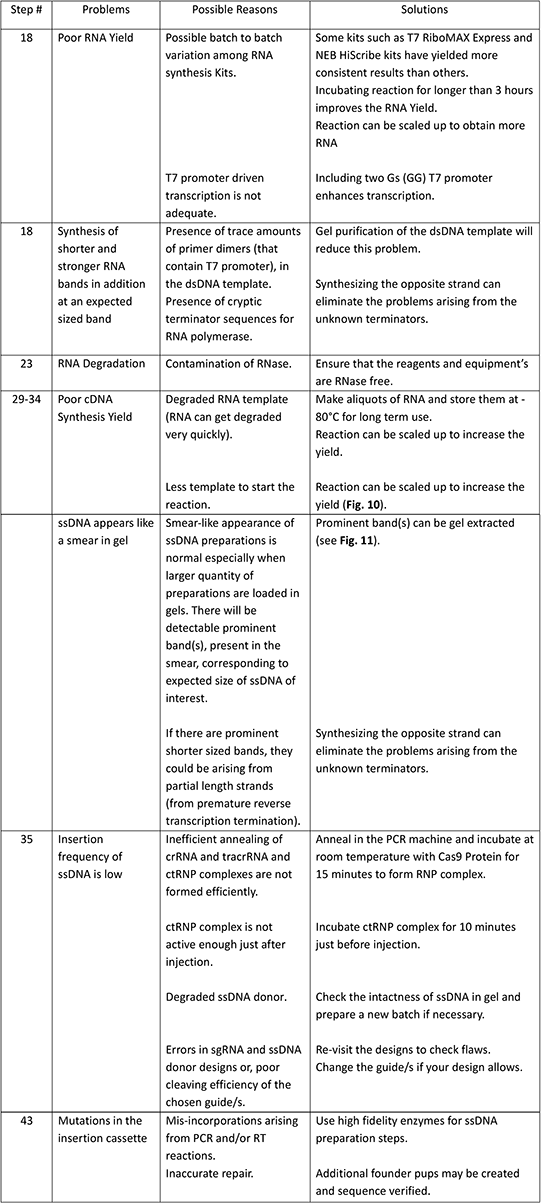

## Anticipated results

*Easi*-CRISPR has the potential to generate various kinds of mouse models including conditional knockout mice (floxed), knock-in mice (to fuse protein expression cassettes with genes), and knock-down mice (artificial microRNA knock-in mice). The method invariably generates intended mouse models in about 50 to 100 of zygotes injected. The method has been successful for over a dozen loci thus far, and the experiments performed in at least three independent laboratories. Described below are the anticipated results discussed under each of the major experimental steps. A separate table is included regarding troubleshooting of experimental steps, which are also briefly discussed in this section.

### Guide search

In the case of floxing and knock-down model designs (that require insertion of artificial microRNA cassettes), it should be easy to find suitable guides because of the flexibility of insertion locations in those introns. Because *LoxP* sites/microRNA cassettes can be placed anywhere within a region that was originally narrowed for searching guides, any guide with a good score and with the lowest or no potential off target cleavage can be chosen for the design. For the knock-in designs (requiring insertion of protein coding fusion cassettes with the gene sequences), the best guide will be that which cleaves exactly at the location where the new sequence needs to be inserted. If such a guide is not available, the next nearest cleaving guide should be chosen (see **Figure 2** for examples of hypothetical guides and their locations with respect to the desired site of insertion). In some cases, we have chosen guides up to about 13 bases away from the insertion site, but have noticed imprecise insertions in some founders (deletion of the 13 nucleotides). If the chosen is farther from the desired insertion site, more founders may need to be generated to obtain correctly inserted founders. Irrespective of the guide cleaving location, the inside termini of homology arms (that meet the new sequence) will remain the same (as though the cassette were still to be inserted at the desired location), while the lengths of homology arms are maintained at least 55 bases long by extending their outside termini.

### Preparation of ssDNA

This involves three steps: (a) preparation of dsDNA templates; (b) preparation of RNA from dsDNA; and (c) preparation of ssDNA from RNA. The anticipated results in each of these steps are:

a. Preparation of dsDNA templates. This step (either using a plasmid source or a PCR product) invariably yields more than several μgs of DNA. If some PCR amplifications result in poor yield, a Taq polymerase from a different vendor may be tried, or the reactions can be scaled up to compensate for the amount needed for the next step.
b. Preparation of RNA from dsDNA. Typical yields range between 5-140μg of RNAs, but are dependent on kits used. Even though about 5 μg of RNA is sufficient for the next step, in our experience a total yield of about ~30 μg indicates an optimal quantity for an *in vitro* transcription reaction of 10 μl volume. We have noticed that some batches of kits and RNA purification columns do not yield consistent results. If low yields are obtained, a new batch of this kit or a kit from a different vendor may be tried. A sample gel image depicting variable results is shown in **Figure 8**. In our laboratories, RNA samples are analyzed using regular TAE gels that are routinely used for DNA analysis. It should be noted that molecular weights do not match with those of dsDNA markers, but such gels provide necessary information quickly enough to proceed to the next step.
c. Preparation of ssDNA from RNA. Typical yields, after the final step of gel purification, range from 0.2 to 2 μg of cDNAs, depending on kits and scale of synthesis used. After complete degradation of RNA by RNaseH treatment, the resulting cDNAs are gel-purified, eluted in a micro-injection buffer, and filtrated to avoid clogging the injection needle. Unlike RNA synthesis and purification kits, in our experience cDNA synthesis reagents and kits are less likely to have batch-to-batch or vendor-to-vendor variations. A sample gel showing comparable performance of three different reverse transcription reactions are shown in **Figure 9**. The effects of the total amount of input RNA and incubation periods on the total yield of cDNA is shown in **Figure 10**. These data indicate that about 5μg of input RNA and at least 10 minutes (per the manufacturer’s recommendations) or longer incubation is necessary to obtain optimal ssDNA yields. Note that gel extraction of cDNA is a critical step to remove partially synthesized ssDNAs, and that the presence of incompletely synthesized ssDNA molecules can hamper insertion of full-length molecules. In this step, we load higher amounts of cDNA preparations to reduce the gel volume. When higher amounts are loaded, we often see a smear with one prominent band and another less prominent band migrating just above the prominent band. This is shown in **Figure 11**, where the prominent band and the slower migrating bands are marked as 1 and 2, respectively. Purification of the two bands and running them at lower concentrations reveal that both bands migrate at the same rate (**Fig. 11**). Based on this experiment, we presume that a differential migration of molecules can occur when loaded at a higher concentration; one reason could be that some molecules may migrate at different rates due to the secondary structures in them. Despite the fact that the majority of ssDNA is lost during purification steps, the amount obtained at the end would be sufficient for at least two-to-three sessions of microinjections, given that 50–200 ng of ssDNA is used for one injection session. Finally, the ssDNA preparation can be tested by subjecting an aliquot to S1 nuclease digestion, which digests it completely if the prep only contains ssDNA (**Fig. 12**).

**Figure 8.**
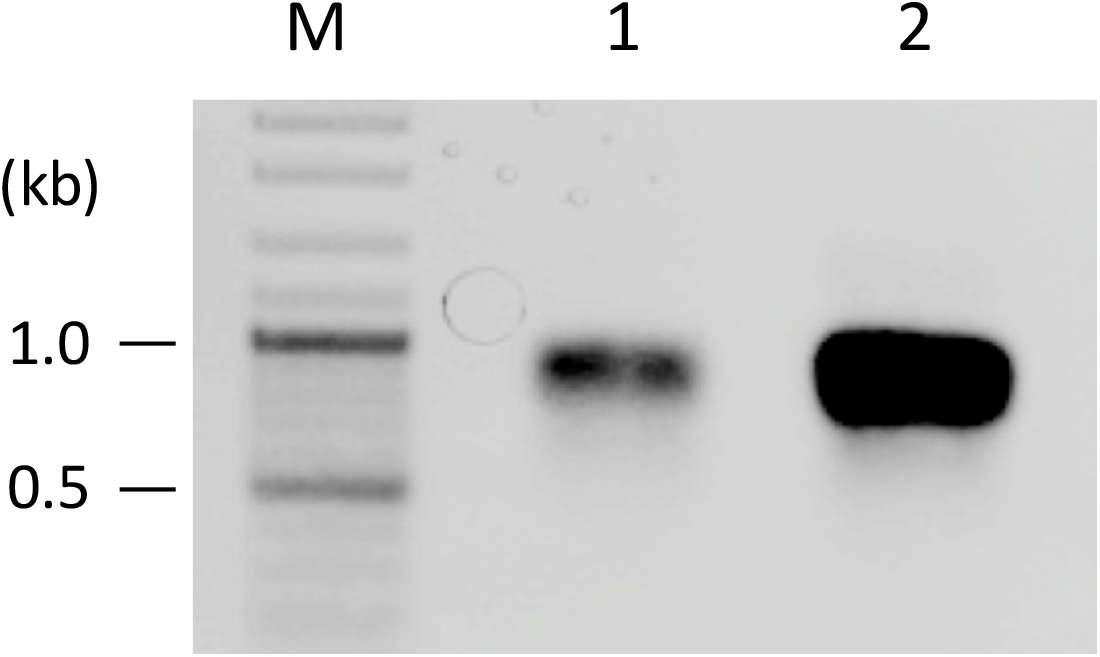
Comparison of RNA yield between kits. 1μg DNA was used for RNA synthesis according to each manufactures protocol. 1 and 2 are RNA synthesized using different RNA synthesis kits from different vendors. After column purification (using MEGAclear), 1% of the reaction volume was loaded. This figure is included to illustrate that kits can vary in their efficiencies (the identity of the products is not an important point here). We have observed a little better performance of a different batch from vendor 1 and somewhat poorer performance of another batch of kit from vendor 2.

**Figure 9.**
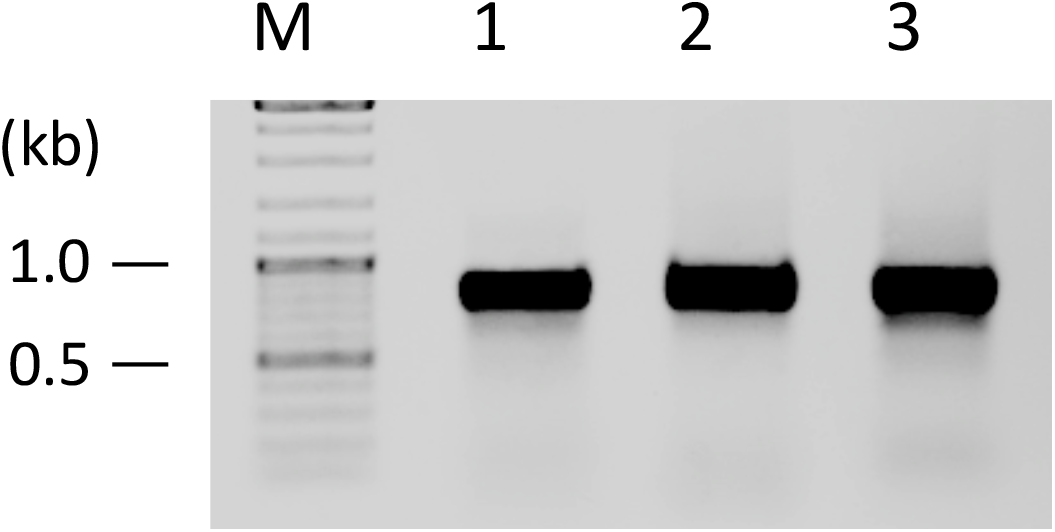
Comparison of cDNA yield among different kits. cDNAs (ssDNA) were synthesized using reverse transcriptases from three different vendors (1 to 3). 1.7μg RNA was used for cDNA synthesis according to each manufactures protocol and 50% of the reaction volumes were loaded.

**Figure 10.**
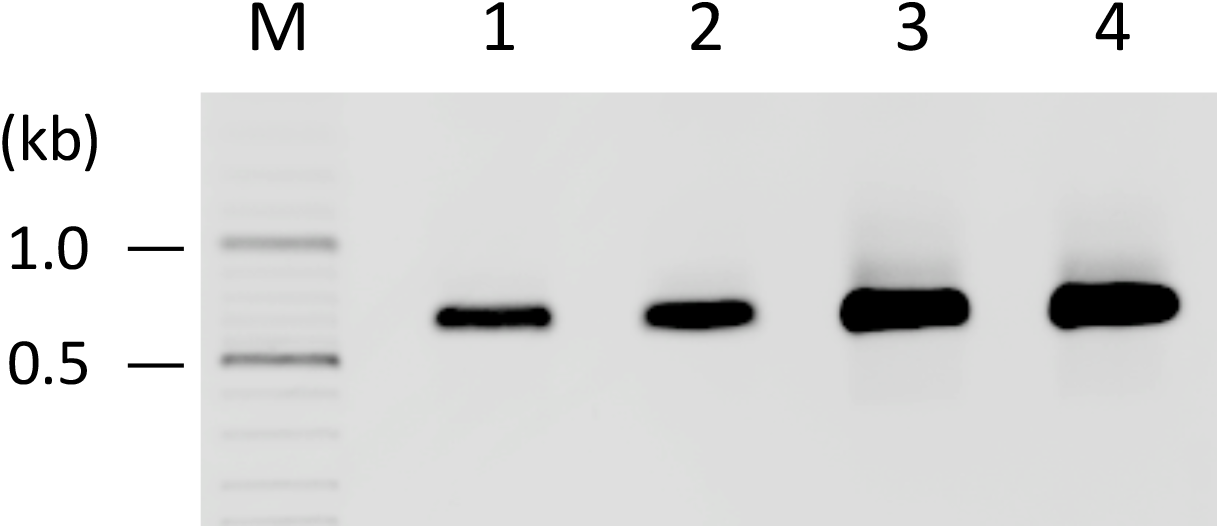
Comparison of cDNA synthesis parameters. 1.7μg (lane 1 and 2) or 5μg (lanes 3 and 4) of RNA was used for cDNA synthesis and the reactions were incubated for 10 minutes (lanes 1 and 3) or 2 hours (lanes 2 and 4). 15% of the reaction volume was loaded.

**Figure 11.**
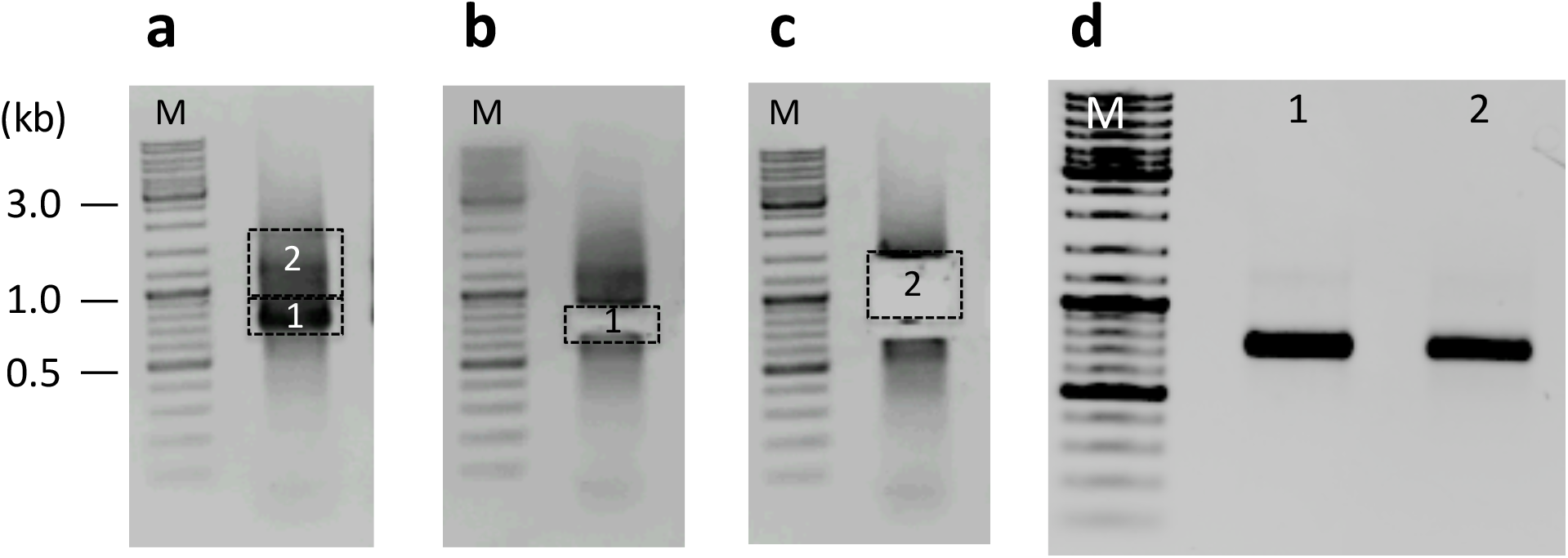
Gel extraction of band/s from the cDNA synthesis preparations. **(a)** Typical smear-like appearance of the cDNA (ssDNA) preparations in an agarose gel showing a prominent band (boxed as 1) and a less prominent band (boxed as 2). **(b and c)** Gel slices excised, purified and loaded in another gel **(d)** Showing both bands migrate similarly. Either of the gel preparations (1 or 2) can be used for microinjection (or both can be combined together if the total is poor).

**Figure 12.**
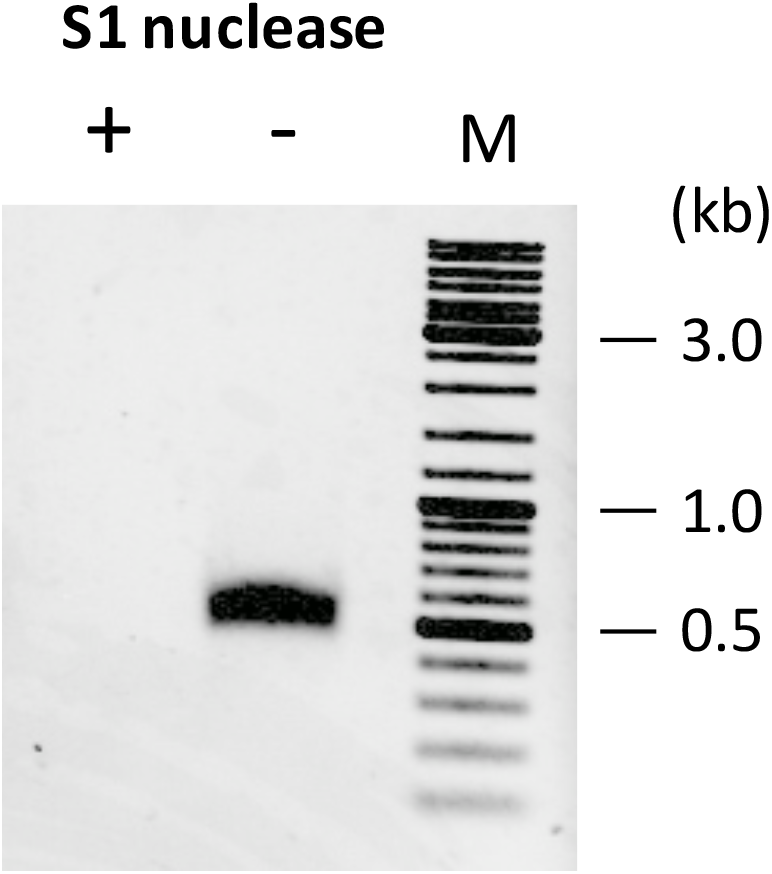
Verification of ssDNA using S1 nuclease. About 150 ng of ~0.5kb cDNA was incubated with or without S1 nuclease at 37°C for 15 minutes.

### Microinjection and transgenesis

Similar to transgenic microinjection experiments, overall birth rates vary from 10 to 30% among *Easi*-CRISPR experiments. However, insertion frequencies among live-born offspring is quite high in Easi-CRISPR, with the majority of projects completed by injecting about 50, or even fewer, zygotes.

### Genotyping

In general, genotyping of CRISPR-generated animal models is very challenging given the potential for mosaicism and for many possible genome editing outcomes, including NHEJ-*indels* and large deletions or insertions. Identifying desired insertion alleles among such mixtures is certainly a difficult task. To ensure identification of correctly targeted alleles, we perform at least three independent sets of PCRs (**Fig. 6**), followed by full sequencing of the targeted alleles. We have noticed higher rates of imprecise or partial insertions in some loci more than others,^8^ which could be attributed to the mechanism of ssDNAs donor-mediated DNA repair, not yet fully understood. Further, some loci may be challenging for PCR genotyping and may require testing of many primers until some are found that work.

### Overall efficiency of *Easi*-CRSPR protocol

Insertion frequencies of ssDNA cassettes is typically 25% to 100% among live-born pups. Cleaving efficiency of the guides could be one of the factors affecting insertion frequencies, and thus some loci are more efficient than others.

## AUTHOR CONTRIBUTIONS

M.O conceived the idea of using long ssDNAs as donors in genome editing, which was further tested and improved upon by the other three authors. All authors contributed equally in writing the manuscript.

## ACKNOWLEDGMENTS

This work was supported in part by Grant-in-Aid for Young Scientists (B) (16K18821) from the Ministry of Education, Culture, Sports, Science and Technology (MEXT) to H.M, and by 2014 Tokai University School of Medicine Research Aid, MEXT-Supported Program for the Strategic Research Foundation at Private Universities 2015–2019 (PI: Yutaka Inagaki), Research and Study Project of Tokai University General Research Organization, 2016-2017 Tokai University School of Medicine Project Research to M.O and by an Institutional Development Award (PI: Shelley Smith) P20GM103471 (to C.B.G and R.M.Q). We also gratefully acknowledge the contribution of the staff of the Support Center for Medical Research and Education, Tokai University, for sequencing and microinjection.

## SUPPLEMENTAL INFORMATION

Supplemental information includes transgenesis experimental steps, which can be found with the article online.

## Supplemental Text: Mouse transgenesis experiments

These experiments follow well-established standard protocols of mouse transgenesis, typically performed at specialized core facility laboratories. Such protocols have been described in detail elsewhere^23,24,29^

### REAGENTS FOR MOUSE TRANSGENSIS EXPERIMENTS

- Pregnant mare serum gonadotropin (PMSG) and Human chorionic gonadotropin (HCG) (National Hormone and Peptide Program (Harbor–UCLA Medical Center, Torrance, CA). Hormones are supplied as lyophilized vials of 2000 IUs. Reconstitute one vial in 2 ml PBS (Millipore, cat. no. BSS-1006-B). Aliquot this 20X stock solution (100 IU/100μl) into 100 μl vials and store at −80°C. For injection, thaw one vial and dilute to 2 ml with PBS to get a final concentration of 5 IU/0.1 ml. Each animal is administered 0.1 ml of this solution
- M2 media for embryo handling (Millipore, cat. no. MR-015-D)
- Hyaluronidase (Millipore, cat. no. MR-051-F)
- KSOM + AA for embryo incubation (Millipore, cat. no. MR-106-D)
- Light mineral oil (Millipore, cat. no. ES-005-C)
- Falcon tissue culture dish 35 × 10 mm (cat. no. 353001)
- Falcon tissue culture dish 60 × 15mm (cat. no. 353002)
- Falcon IVF dish (cat. no. 353653)
- Glass capillaries (4 in/1 mm) (World Precision Instruments, cat. no. TW100F-4)
- Micro Fill 28 gauge/97mm long. (World Precision Instrument Inc. Item # MF28G)
- Flexipet oocyte/embryo pipettes (Cook Medical K-FPIP-1130-10BS-5)
- Tuberculin syringes (NORM-JECT 4010-200V0)
- Holding pipettes. (Humagen MPH-SM-20)
- Chamber slide (Lab-Tek, cat. no. 177372)
- Embryo Max Microinjection buffer (cat.no. MR-095-10F)

### MICE

- **Donor females**: Purchase three week old C57BL/6N females from Charles River Laboratories (Wilmington, MA)
- **Stud males**: Purchase C57BL/6N males from Charles River Laboratories (Wilmington, MA)
- **Pseudopregnant recipients**: Purchase Crl: CD-1(ICR) female mice at 5–6 weeks of age from Charles River Laboratories (Wilmington, MA)
- **Vasectomized males**: Purchase 5–6 week old CD-1 mice from Charles River Laboratories (Wilmington, MA) and perform vasectomies as described previously^30^. **! CAUTION** Experimental procedures involving animals should be carried out according to relevant institutional and governmental regulations.

### EQUIPMENT

- Glass pipette puller (Sutter Instrument Co. model #P97), outfitted with a 2.5mm X 2.5mm Box filament (cat. no. FB255B)
- Glass Capillaries World Precision Instrument Item # TW100F-4 w/Filament 1.0mm 4in. Sarasota FL. USA
- Nikon Eclipse TE 2000-E w/ DIC equipped with Narishige IM 300 microinjector and NT-88-V3 micromanipulators
- Heating glass. Controller (CU-301) HG-T-Z002. Live Cell Instruments
- Leica DM IRB equipped with Narishige IM 300 microinjector and Leica manual manipulators
- Leica MZ 9.5

▪ Condenser lens: PLAN 0.5x, model #10 446 157
▪ Base: Model #10 445 367
▪ Tilt head
▪ Heating glass (Live Cell Instrument, cat. no. HG-T-Z002) with temperature controller (Live Cell Instrument, cat. no. CU-301)
- Heraeus Hera cell 150 Tri-gas incubator equipped with Coda inline filters
- Nunc Lab-Tek Chamber Slide System (Lab-Tek, cat. no. 177372)
- Standard surgical equipment such as scissors, fine forceps, suturing material, anesthesia chambers, etc.
- Large slide warmer (Spectrum Scientifics, cat. no. 3875)
- Mouth pipetting apparatus
- Heating glass (Live Cell Instrument, cat. no. HG-T-Z002) with temperature controller (Live Cell Instrument, CU-301)
- Microinjection scope: Leica DM IRB, equipped with Narishige IM 300 microinjector, and Leica manual manipulators

▪ Eyepiece: HC PLAN 10x/22 w/tilt, model #11 507 804
▪ Condenser lens:.30 S70
▪ Objectives:

- C PLAN 4X/.10, model #11 506 074
- N PLAN L20X/0.40 CORR, model #11 506 057
- N PLAN L40X/0.55 CORR, model #11 506 059

### Protocol steps of mouse transgenesis (these steps are performed between the steps 41 and 42 of the *Easi*-CRISPR protocol). **● TIMING** 5-6h

1. Isolation of embryos and microinjection steps are very similar to the protocols followed for mouse genome editing using CRISPR/Cas9^23^.
2. Prepare a slide using a Lab-Tek chamber by making two side-by-side 150 μl drops of M2 media. Flatten the drops into discs with a pipette tip to minimize their height. Overlay the drops with ~1 ml of mineral oil. **▲ CRITICAL STEP** Maintain the temperature at 37°C on a slide warmer.
3. Inject about 20–30 zygotes per batch. All zygotes, in each batch, must be injected within 10 minutes, thus the number of zygotes taken per batch depends on the efficiency of the injector. A beginner may start with as few as 4 to 6 per batch; an experienced technician can inject as many as 50 zygotes in 10 minutes.
4. Perform microinjection.

a. Maintain positive pressure on the injection needle at all times.
b. Penetrate the needle into zona pellucida. Bring the needle to the closest pronucleus.
c. A slight swelling of the pronucleus may be seen once the membrane is penetrated. Otherwise, press the injection foot pedal to observe a slight swelling of the pronucleus.
d. Carefully withdraw the capillary from the zygote.
5. After all zygotes are injected, use the mouth pipetting apparatus to collect and transfer zygotes to the embryo transfer dish. **? TROUBLESHOOTING** **▲ CRITICAL STEP** Discard lysed zygotes.
6. Incubate surviving zygotes at 37°C in KSOM until embryo transfer (usually within the next 1–2 hours).
7. Transfer of injected embryos into pseudo-pregnant mice.

a. Obtain pseudo-pregnant mice by mating 8–12 week old CD-1 females to vasectomized CD-1 males.
b. On the morning of injection day, use plug-positive females for oviduct transfers. **▲ CRITICAL STEP** Typically, 10–20 CD-1 females are bred in each session to obtain an average of 4–8 plugged females.
c. Transfer viable manipulated embryos to the oviducts of pseudo-pregnant foster mothers, following established surgical procedures as previously described^30^. About 15–25 injected embryos are transferred per female. The optimal number of embryos transferred is 18 total per female (9 per side). **? TROUBLESHOOTING**

### Troubleshooting (supplementary procedures)

| STEP | Problem | Possible Reason | Solution |
| --- | --- | --- | --- |
| Step 5 | Embryo lysis during microinjection | Clogging of Injection Needles | Centrifuging and Filtration of injection can prevent clogging of needles. |
| Step 7c | Loss of Embryos and low birth rate | Higher concentration of ssDNA, or some component in the injection mix may be toxic.<br>Disruption of gene of interest if the gene is involved in embryonic development (due to introduction of *indel* instead of targeted insertion), or if the guide has an off target effect on another gene necessary for embryonic viability | Reduce the concentration of ssDNA in the injection mix (5ng/ul or less), purchase a new batch of crRNA if the toxicity persists.<br>Change the guide(s), or the strategy, if the project allows |

## REFERENCES

1. Gurumurthy, C. B. et al. Validation of simple sequence length polymorphism regions of commonly used mouse strains for marker assisted speed congenics screening. Int. J. Genomics 2015, 735845 (2015).

2. Gurumurthy, C. B. et al. CRISPR: a versatile tool for both forward and reverse genetics research. Hum. Genet. 135, 971–976 (2016).

3. Quadros, R. M., Harms, D. W., Ohtsuka, M. & Gurumurthy, C. B. Insertion of sequences at the original provirus integration site of mouse ROSA26 locus using the CRISPR/Cas9 system. FEBS Open Bio 5, 191–197 (2015).

4. Inui, M. et al. Rapid generation of mouse models with defined point mutations by the CRISPR/Cas9 system. Sci. Rep. 4, (2014).

5. Horii, T. & Hatada, I. Challenges to increasing targeting efficiency in genome engineering. J. Reprod. Dev. 62, 7–9 (2016).

6. Yang, H. et al. One-step generation of mice carrying reporter and conditional alleles by CRISPR/Cas-mediated genome engineering. Cell 154, 1370–1379 (2013).

7. Ma, Y. et al. Generating rats with conditional alleles using CRISPR/Cas9. Cell Res. 24, 122–125 (2014).

8. Miura, H., Gurumurthy, C. B., Sato, T., Sato, M. & Ohtsuka, M. CRISPR/Cas9-based generation of knockdown mice by intronic insertion of artificial microRNA using longer single-stranded DNA. Sci. Rep. 5, 12799 (2015).

9. Quadros, R. M. et al. Easi-CRISPR: a robust method for one-step generation of mice carrying conditional and insertion alleles using long ssDNA donors and CRISPR ribonucleoproteins. Genome Biol. 18, (2017).

10. Jacobi, A. M. et al. Simplified CRISPR tools for efficient genome editing and streamlined protocols for their delivery into mammalian cells and mouse zygotes. Methods (2017). doi:10.1016/j.ymeth.2017.03.021

11. Suda, T., Gao, X., Stolz, D. B. & Liu, D. Structural impact of hydrodynamic injection on mouse liver. Gene Ther. (2006). doi:10.1038/sj.gt.3302865

12. Takahashi, G. et al. GONAD: Genome-editing via Oviductal Nucleic Acids Delivery system: a novel microinjection independent genome engineering method in mice. Sci. Rep. 5, 11406 (2015).

13. Cong, L. et al. Multiplex Genome Engineering Using CRISPR/Cas Systems. Science 339, 819–823 (2013).

14. Mali, P. et al. RNA-Guided Human Genome Engineering via Cas9. Science 339, 823–826 (2013).

15. Skarnes, W. C. Is mouse embryonic stem cell technology obsolete? Genome Biol. 16, (2015).

16. Cohen, J. ‘Any idiot can do it.’ Genome editor CRISPR could put mutant mice in everyone’s reach. Science (2016). doi:10.1126/science.aal0334

17. Maruyama, T. et al. Increasing the efficiency of precise genome editing with CRISPR-Cas9 by inhibition of nonhomologous end joining. Nat. Biotechnol. 33, 538–542 (2015).

18. Nakao, H. et al. A possible aid in targeted insertion of large DNA elements by CRISPR/Cas in mouse zygotes. Genes. N. Y. N 2000 54, 65–77 (2016).

19. Wang, B. et al. Highly efficient CRISPR/HDR-mediated knock-in for mouse embryonic stem cells and zygotes. BioTechniques 59, (2015).

20. Aida, T. et al. Gene cassette knock-in in mammalian cells and zygotes by enhanced MMEJ. BMC Genomics 17, (2016).

21. Bishop, K. A. et al. CRISPR/Cas9-Mediated Insertion of loxP Sites in the Mouse Dock7 Gene Provides an Effective Alternative to Use of Targeted Embryonic Stem Cells. G3 Bethesda Md 6, 2051–2061 (2016).

22. Lee, A. Y. & Lloyd, K. C. K. Conditional targeting of Ispd using paired Cas9 nickase and a single DNA template in mice. FEBS Open Bio 4, 637–642 (2014).

23. Harms, D. W. et al. Mouse Genome Editing Using the CRISPR/Cas System. Curr. Protoc. Hum. Genet. Editor. Board Jonathan Haines Al 83, 15.7.1–15.7.27 (2014).

24. Yang, H., Wang, H. & Jaenisch, R. Generating genetically modified mice using CRISPR/Cas-mediated genome engineering. Nat. Protoc. 9, 1956–1968 (2014).

25. Yoshimi, K. et al. ssODN-mediated knock-in with CRISPR-Cas for large genomic regions in zygotes. Nat. Commun. 7, 10431 (2016).

26. Aida, T. et al. Cloning-free CRISPR/Cas system facilitates functional cassette knock-in in mice. Genome Biol. 16, (2015).

27. Ellefson, J. W. et al. Synthetic evolutionary origin of a proofreading reverse transcriptase. Science 352, 1590–1593 (2016).

28. Hendel, A. et al. Chemically modified guide RNAs enhance CRISPR-Cas genome editing in human primary cells. Nat. Biotechnol. 33, 985–989 (2015).

29. Schilit, S. L. P., Ohtsuka, M., Quadros, R. M. & Gurumurthy, C. B. in Current Protocols in Human Genetics (eds. Haines, J. L. et al.) 15.10.1-15.10.28 (John Wiley & Sons, Inc., 2016).

30. Behringer, R., Gertsenstein, M., Nagy, K. V. & Nagy, A. Manipulating the mouse embryo: a laboratory manual. (2014).

